# Barcoding of small extracellular vesicles with CRISPR-gRNA enables comprehensive, subpopulation-specific analysis of their biogenesis/release regulators

**DOI:** 10.1101/2023.09.28.559700

**Authors:** Koki Kunitake, Tadahaya Mizuno, Kazuki Hattori, Chitose Oneyama, Mako Kamiya, Sadao Ota, Yasuteru Urano, Ryosuke Kojima

**Affiliations:** Graduate School of Pharmaceutical Sciences, The University of Tokyo, 113-0033 Tokyo, Japan; Research Center for Advanced Science and Technology, The University of Tokyo, 153-8904 Tokyo, Japan; Division of Cancer Cell Regulation, Aichi Cancer Center Research Institute, 464-8681 Nagoya, Japan; Department of Life Science and Technology, Tokyo Institute of Technology, 226-8501 Kanagawa, Japan; Graduate School of Medicine, The University of Tokyo, Tokyo 113-0033, Japan; PRESTO, Japan Science and Technology Agency, Kawaguchi, Saitama, 332-0012, Japan; FOREST, Japan Science and Technology Agency, Kawaguchi, Saitama, 332-0012, Japan

## Abstract

Small extracellular vesicles (sEVs) are important intercellular information transmitters in various biological contexts, but their release processes remain poorly understood. Herein, we describe a high-throughput assay platform, <u>C</u>RISPR-assisted individually <u>b</u>arcoded s<u>E</u>V-based release <u>r</u>egulator (CIBER) screening, for identifying key players in sEV release. CIBER screening employs sEVs barcoded with CRISPR-gRNA through the interaction of gRNA and dead Cas9 fused with an sEV marker. Barcode quantification enables the estimation of the sEV amount released from each cell in a massively parallel manner. Barcoding sEVs with different sEV markers in a CRISPR pooled-screening format allows genome-wide exploration of sEV release regulators in a subpopulation-specific manner, successfully identifying previously unknown sEV release regulators and uncovering the exosomal/ectosomal nature of CD63^+^/CD9^+^ sEVs, respectively, as well as the synchronization of CD9^+^ sEV release with the cell cycle. CIBER should be a valuable tool for detailed studies on the biogenesis, release, and heterogeneity of sEVs.

---

Membrane-enclosed extracellular vesicles (EVs) released by cells are typically classified into several subgroups depending on their size or origin, including small EVs (sEVs, 30-200 nm in diameter), medium/large EVs (EVs larger than sEVs), exosomes (multivesicular body (MVB) derived), and ectosomes (plasma-membrane derived, also known as microvesicles)^1^. Among them, sEVs containing various biomolecules are important mediators of cell-to-cell communication in both physiological and pathological contexts, including cancer metastasis^2,3^. These facts highlight the potential of sEV biogenesis and release processes (“release” processes hereafter) as novel therapeutic targets^3,4^. Furthermore, sEVs are also attracting attention as highly biocompatible delivery vesicles^3,5^, and therefore methods to control/enhance their production are of great interest for biotechnological applications. Despite the importance of sEV release processes, a comprehensive understanding of their regulation has remained elusive for several reasons^6–9^. Firstly, multiple biological molecules are involved in interconnected pathways, which are difficult to analyze with conventional low-throughput assays using small-molecule inhibitors or siRNAs in separate wells (Fig.1a, upper). Secondly, many of the regulators of sEV release also affect cellular activities, including viability, which hampers high-throughput identification of factors controlling sEV release. Thirdly, sEVs are heterogeneous, and the release mechanisms of different subpopulations of sEVs are hard to differentiate.

**Figure 1.**
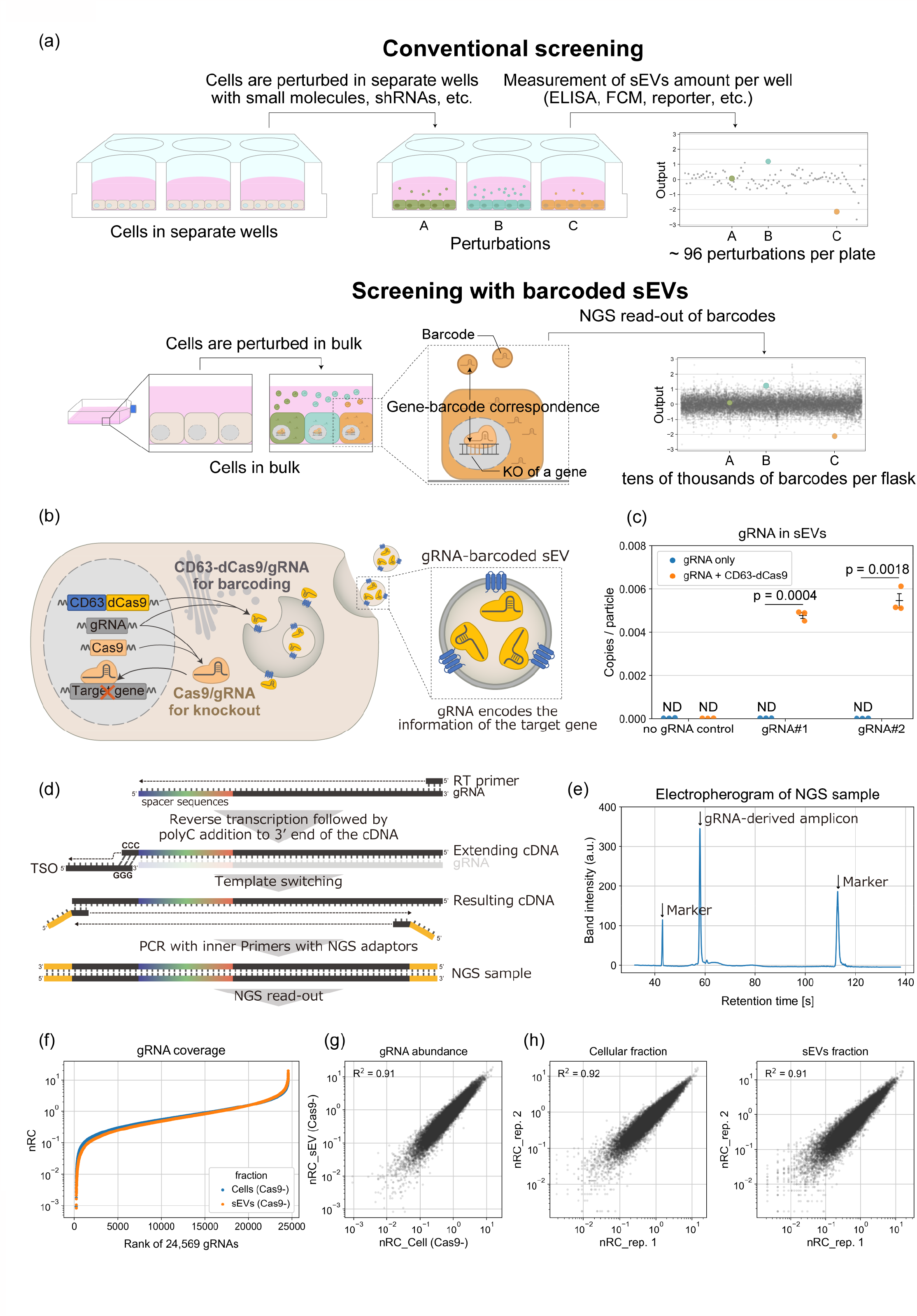
Concept and creation of gRNA-barcoded sEVs for high-throughput analyses of their release regulators (a) Schematic illustration of the use of “barcoded” sEVs for analyses of their release regulators. Conventional biological assays (e.g., by treating cells with a small-molecule inhibitor or siRNA in separate wells and measuring the amount of sEVs released in each well) offer only rather low throughput. On the other hand, sEVs barcoded with CRISPR guide RNA (gRNA) allow for the identification of sEV release regulators in a pooled manner via read-out of gRNA composition by NGS. Another advantage of this pooled screening would be that it is possible to exclude the effect of sEV re-uptake on the amount of sEVs by randomizing the cellular environment (i.e., in a setting that employs separate wells, downregulation/upregulation of sEV re-uptake might be misinterpreted as upregulated/downregulated sEV release, respectively). (b) Schematic illustration of the generation of sEVs barcoded with gRNA. By expressing a fusion protein of CD63 (sEV marker) and dead Cas9 (dCas9, RNA binding protein for gRNA) together with Cas9 in the sEV-producing cells, gRNA used for gene knockout can be actively encapsulated in sEVs. (c) gRNA abundance in sEVs isolated from culture media of HEK293T or HEK293T cells expressing CD63-dCas9 stably transduced with gRNA using lentivirus. The copy numbers of gRNAs were determined by qPCR (using gRNA-encoding plasmid as a standard) and divided by the particle number measured by Nanosight to calculate copies/particle. The Ct values of qPCR for gRNA#1 and #2 without CD63-dCas9 were not significantly different from the no-gRNA control (the observed Ct should represent non-specific amplification), so the copy numbers are shown as ND (not determined). p: one sample t-test compared to 0. Error bars represent ± SEM of biological replicates (n = 3). (d) Developed method for spacer amplification for next generation sequencing (NGS). Common adapter sequences are incorporated during first-strand cDNA synthesis from a mixture of gRNAs with various spacer sequences using a reverse transcriptase having template-switching capability. Then, the cDNA is amplified by nested PCR with a common pair of primers for NGS analysis. (e) A Bioanalyzer electropherogram of an NGS sample prepared from sEV RNA as shown in Fig 1c. (f) Coverage of gRNAs. Cells expressing CD63-dCas9 were transduced with a library of 24,569 gRNAs using lentivirus (DTKP library, see Supplementary Fig. 6). RNAs extracted from cells and sEVs released from them were processed for NGS read-out as shown in Fig. 1c. Read counts for every single gRNA were divided by the total sample count, then multiplied by the number of gRNAs in the library (24,569) to calculate the normalized read count (nRC), such that the mean value becomes 1. Data are shown as mean nRC from two biological replicates. (g) Correlation between nRCs of 24,569 gRNAs in sEVs and cellular fractions from 2 replicate cultures of Cas9-cells. (h) Reproducibility of nRCs of 24,569 gRNAs from cellular and sEVs fractions measured by NGS.

Here we report a novel pooled screening system that overcomes these limitations and its application to identify key players of sEV release processes. We actively incorporated guide RNA (gRNA) for Cas9 into sEVs through the interaction of gRNA and dead Cas9 (dCas9) fused with an sEV marker in a pooled CRISPR screening format. This allows sEV-loaded gRNA to work as a “barcode” linking each sEV to the perturbation of gene expression in its originating cell. Quantification of the composition of barcode gRNA in both sEVs and cells allows high-throughput, genome-wide exploration of genes involved in sEV release while canceling out the effects on cellular activities (e.g., proliferation, barcode transcription). We call this assay platform CRISPR-assisted individually barcoded sEV-based release regulator (CIBER) screening. CIBER screening using multiple sEV markers in combination with bioinformatic analyses revealed both known and previously unknown factors controlling sEV release processes, uncovering different effects of V-type ATPases, mitochondrial electron transport, and the cell cycle on the release of CD63^+^ and CD9^+^ sEVs. We believe this work provides a basis for detailed studies on the biogenesis, release, and heterogeneity of sEVs. We discuss the potential of this sEV-barcoding platform for various future applications.

## Results

### Concept and design of a library of sEVs barcoded with CRISPR-Cas9 gRNA

In CRISPR pooled screening, Cas9-expressing cells are transduced with a library of gRNAs to perturb gene expression. These genome-incorporated gRNAs in turn enable estimation of the numbers of cells bearing different gene knockouts via read-out of the gRNA by next-generation sequencing (NGS) (Supplementary Fig. 1)^10^. Analogously, we hypothesized that it would be possible to estimate the numbers of sEVs if we could efficiently load sEVs with gRNA transcribed in their originating cells, thereby enabling identification of regulators of sEV release in a pooled, massively parallel manner (Fig.1a, lower). In this setting, each transcribed gRNA will encode information regarding the perturbed gene in its originating cell as a barcode, and quantification of the gRNA barcodes in sEVs by next-generation sequencing (NGS) should allow quantification of sEVs released from each cell.

It has been suggested that stochastic packaging of biomolecules into sEVs is a rather rare event^11^. Therefore, simple overexpression of gRNA inside the cells would be insufficient to create high-quality gRNA-barcoded sEVs. On the other hand, we and several other researchers have reported that specific RNAs can be actively loaded into sEVs by synthetic interaction with an RNA-binding protein (RBP) fused with an sEV marker protein (e.g. CD63)^12–14^. Building on this work, we expected that gRNA could be efficiently loaded into sEVs by using dCas9 as a strong RBP for gRNA (Fig. 1b). Indeed, when we infected HEK293T cells expressing CD63-dCas9 with lentivirus encoding single gRNAs, approximately 1 copy of gRNA per 200 sEVs was detected by qPCR, while the gRNA in sEVs was nearly undetectable in the absence of CD63-dCas9 (Fig. 1c).

Next, we developed a new strategy to amplify gRNA spacers directly from a mixture of transcribed gRNAs (Fig. 1d). Unlike normal CRISPR screening that decodes gRNA spacers in the genome, this new approach is essential for the parallel analysis of multiple gRNAs in sEVs by next-generation sequencing (NGS). Considering that the spacer is located at the 5’-end of gRNA, we adopted SMART technology, which adds a common sequence at the 5’-end during reverse transcription by template switching^15^. By employing nested PCR, we could selectively amplify the gRNA spacers from RNAs in sEV with a common pair of primers (Fig. 1e).

We then infected HEK293T cells expressing CD63-dCas9 with a library of 24,569 gRNAs^16^. With the developed spacer amplification method, we succeeded in detecting about 99% of spacer sequences by NGS with a low skew ratio in both cells and sEVs (Fig. 1f, Supplementary data 1). Importantly, when we compared the abundance of each gRNA spacer in the cellular and sEV fractions, a linear relationship was observed (Fig. 1g), indicating that each gRNA was loaded uniformly into sEVs regardless of its spacer sequence. It is noteworthy that 500 cells / gRNA was sufficient to reproducibly achieve such high barcoding-decoding performance (Fig. 1h), because this cell number is comparable to that utilized in normal pooled CRISPR screening^10^. We also note that the selection of dCas9 as the strong RBP for gRNA was important because the sEV barcoding efficacy was much lower when a bacteriophage MS2 coat protein was used as the RBP for gRNA bearing MS2-binding motifs^17^ (Supplementary Fig. 2). After confirming that the expression of CD63-dCas9 does not change sEV size (Supplementary Fig. 3), that the barcoding performance was consistently high when Cas9 was additionally expressed (Supplementary Fig. 4, Supplementary Data 1,2), and that the co-expression of CD63-dCas9 does not competitively inhibit gRNA-driven gene knockout in Cas9-expressing cells (Supplementary Fig. 5), we decided to apply this dCas9-based sEV barcoding strategy for downstream screening.

### Quantitative evaluation of the effect of Cas9-induced perturbation on sEVs release

HEK293T cells expressing CD63-dCas9 with/without Cas9 (hereafter called Cas9+ and Cas9-cells in this section) were infected with lentivirus encoding CRISPR knockout sub-pool gRNA libraries^16^ listed in Supplementary Fig. 6 (covering 10527 genes in total) and gRNAs were extracted from both sEV and cellular fractions (Supplementary Fig. 6, Supplementary Data 1,2). Although exploration of sEV release regulators could in principle be performed by comparing the gRNA abundances in sEV fractions from Cas9+ and Cas9-cells, it is important to consider that the amount of each gRNA in the sEV fraction would also be affected by the change in the cellular fraction due to the gene knockout (e.g., changes of proliferation rate, barcode transcription, etc.; Fig 2a). Indeed, when we checked the log_2_(fold change (FC)) of the abundance of each gRNA between Cas9+ and Cas9-samples in sEVs and cells (hereafter referred to as FC_sEVs_ and FC_cells_, respectively), gRNAs targeting POLR3 subunits (regulating the transcription of gRNAs^18^) generally showed similarly low values for both parameters (Supplementary Fig. 7). Furthermore, when we plotted FC_sEVs_ and FC_cells_ of all gRNAs used, a linear relationship was observed (Fig. 2b(1), Supplementary Data 3). These facts indicate that the change of gRNA level in sEVs is highly dependent on the change of cellular gRNA level, and this led us to introduce a score called “release effect” (RE) to find the true sEV release regulators. First, we performed linear regression using FC_sEVs_ and FC_cells_ and defined the ‘RE at the gRNA level’ as a residue with respect to the regression line. Then, the median RE among gRNAs targeting the same gene was adopted as the RE at the gene level, and the z-normalized RE (z-RE) was used as the indicator of the contribution of each gene to sEV release (Fig. 2b). This screening pipeline was designated as **<u>C</u>**RISPR-assisted **<u>I</u>**ndividually **<u>B</u>**arcoded s**<u>E</u>**V-based release **<u>R</u>**egulator (CIBER) screening, in which the z-REs of genes that enhance/suppress sEVs release upon KO should be larger/smaller than 0, respectively.

**Figure 2.**
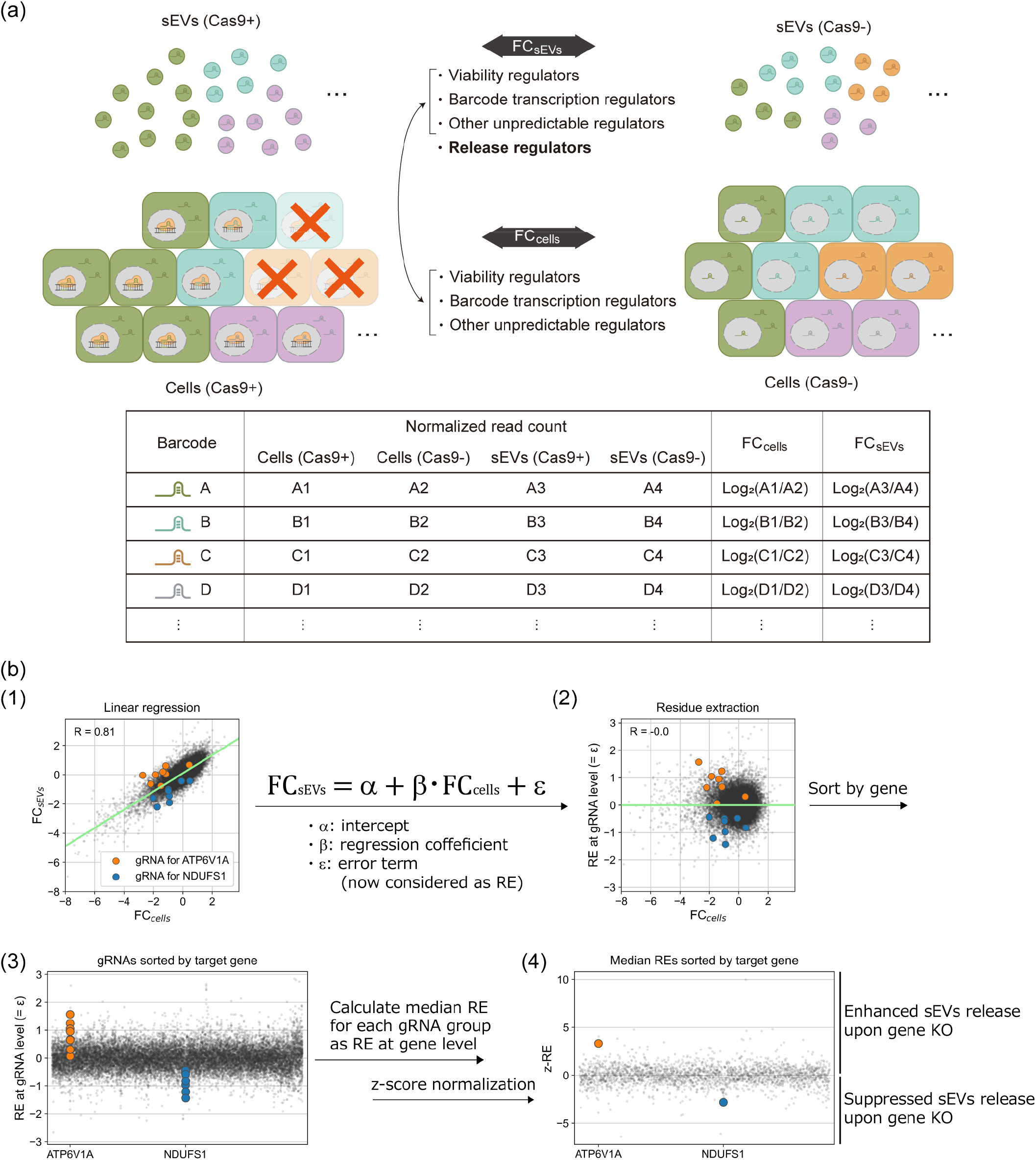
Procedure of CIBER screening. (a) HEK293T CD63-dCas9 cells with/without Cas9 are lentivirally transduced with the gRNA libraries, and the RNA is harvested from both cellular and sEV fractions of these cells. After NGS analysis of the gRNA composition in each sample, FC_sEVs_ and FC_cells_ (FC stands for fold change) of each gRNA are calculated as described. The FC_cells_ of each gRNA reflects the change in the abundance of each gRNA in cells due to the KO of viability regulators, barcode transcription regulators, etc. FC_sEVs_ reflects the effect of sEV release regulators in addition to the factors affecting FC_cells_. Therefore, the actual contribution of each gRNA (and gene) to the change of sEV release can be estimated from both FC_sEVs_ and FC_cells_ by means of the following procedure. (b) Calculation of z-normalized release effect (z-RE) from raw gRNA read counts. (1) Each gRNA was plotted with FC_sEVs_ on the y-axis and FC_cells_ on the x-axis. A linear relationship between FC_cells_ and FC_sEVs_ is observed. The regression line is shown in green (the results for the DTKP library, consisting of 24,569 gRNAs (see Supplementary Fig.6), are shown: R = 0.81). For clarity, gRNAs targeting ATP6V1A and NDUFS1 are highlighted in different colors. (2) Residue values of each gRNA obtained by performing linear regression on (1) are displayed as RE at the gRNA level. (3) gRNAs are sorted according to their target genes. (4) z-RE of each gene is calculated by taking the median value of RE at the gRNA level.

### Results of CD63-dCas9-based CIBER screening

The results of the CD63-dCas9-based CIBER screening (hereafter CD63-CIBER) are shown in Fig. 3a and Supplementary Data 1-4. The genes showing z-RE of >1.65 and <-1.65^19^ are treated as “upper hits” and “lower hits”, respectively. Firstly, we should emphasize that CD63 was detected as one of the top lower hits. This is exactly what we expected, because CD63-targeting gRNAs induce KO of CD63-dCas9 as well, which prevents gRNAs from being loaded into sEVs. Motivated by this confirmation that the positive control gene worked, we validated the screening results with an orthogonal assay using a luciferase reporter CD63-nluc that enables the estimation of CD63^+^ sEV amount by simple luminescence measurement of the cell culture supernatant^12,20^. FASN and PI4KA were among the top lower/upper hits, respectively, and indeed, treatment of the cells with small-molecule inhibitors drastically downregulated/upregulated the CD63-nluc signal, respectively (Fig. 3b, c). These effects were also directly observed by nanoparticle tracking analysis (NTA) with native sEVs (Fig. 3d, e, Supplementary Fig. 8), confirming that CIBER screening can identify new regulators of sEV release. The multiple hit genes highlighted in Fig. 3a were also validated by siRNA-based assays (Fig. 3f, Supplementary Fig. 9).

**Figure 3.**
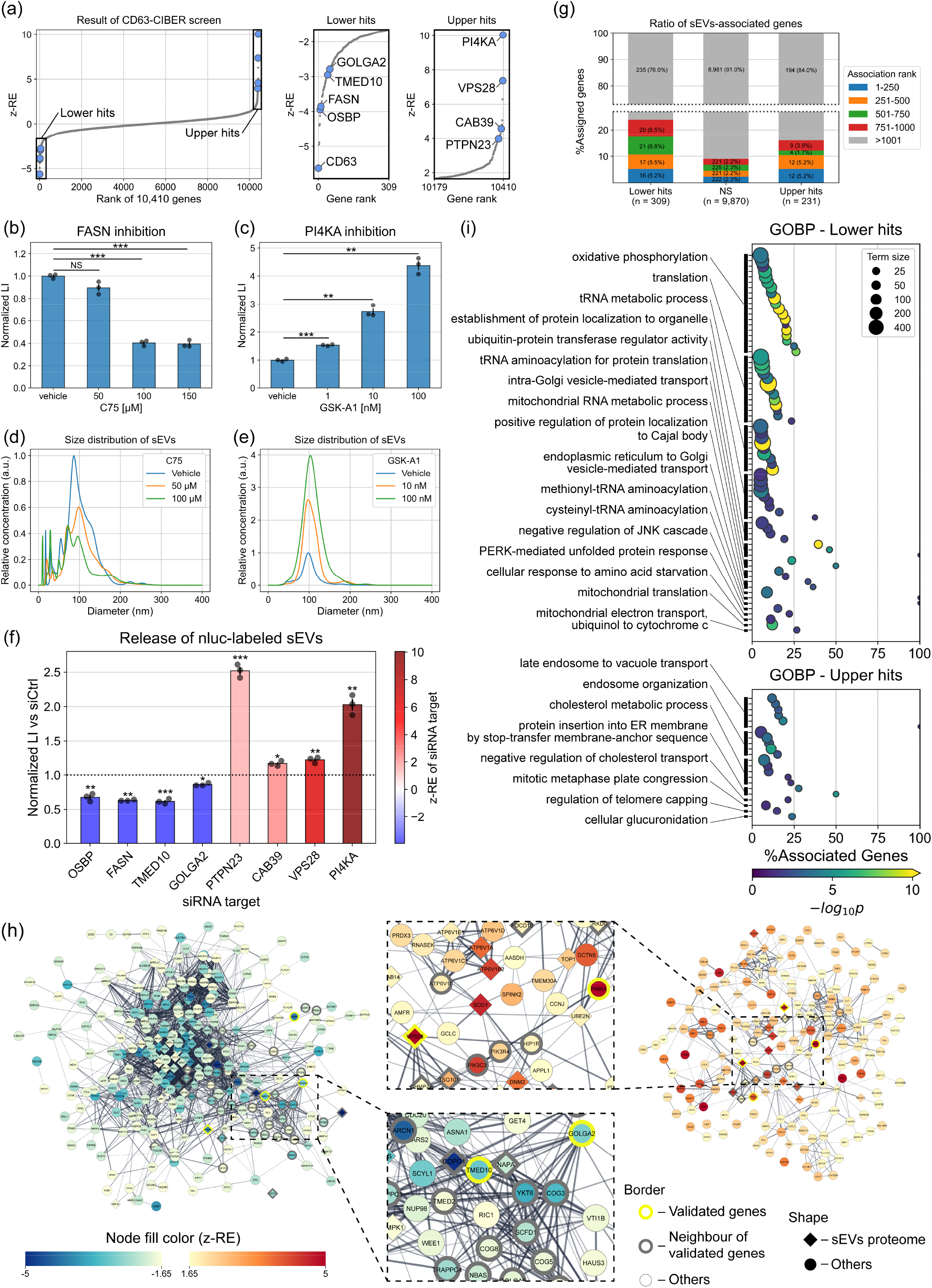
The results of the CD63-dCas9-based CIBER screening. (a) z-RE of 10,410 genes (left panel). The genes showing z-RE larger than 1.65 and lower than -1.65 are considered as “upper hits” and “lower hits”, respectively, and are displayed in magnified panels (center and right). Individual z-RE values for gRNAs and genes are listed in Supplementary Data 3 and 4. (b, c) Validation of the effects of PI4KA and FASN by means of CD63-nanoluc (nluc) reporter assay. HEK293T cells stably expressing CD63-nluc were treated with various concentrations of C75, a FASN inhibitor (b) or GSK-A1, a selective PI4KA inhibitor (c). After each treatment, culture media (CM) were collected and the nanoluc luminescence was measured after stepwise centrifugation to remove the signals from cells, cell debris, etc. The luminescence intensity of each sample was normalized by the protein amount in the cellular fraction measured by BCA assay to cancel out the effect of cell viability. (d, e) Size distribution of sEVs in CM of HEK293T treated with C75 (d) or GSK-A1 (e). CM was cleared by stepwise centrifugation before the NTA measurement. Each result is the average of 3 sequential measurements of each sample. (f) CD63-nluc reporter assay for hit genes by siRNAs. Cells are transfected with siRNA targeting one of the hit genes and processed as shown in Fig. 3b, c. (g) Percent of sEVs-associated genes for Lower hits (not significant, NS) and Upper hits samples. sEVs-associated genes were compiled from the Vesiclepedia database. (h) Protein-protein interaction (PPI) network among lower hits (left) and upper hits (right) analyzed by StringApp in Cytoscape. Each node represents a gene product, and the node color corresponds to the z-RE. Only nodes connected to at least one other node are shown. The density of the edge is proportional to the strength of the PPI evidence (significance score). The border of the nodes of genes validated in Fig. 3f is shown in thick yellow; nodes directly connected to validated genes are shown in thick black. The nodes of proteins found in the sEVs proteome (Fig. S11) are diamond-shaped while others are circular. (i) Results of gene ontology biological process (GOBP) enrichment analysis. The x-axis represents the gene ratio, which refers to the ratio of lower or upper hits genes to all gene numbers annotated to the term. The circle size indicates the total number of genes annotated to the term. The circle color indicates the -log_10_p-value by Fisher’s exact test adjusted by Holm correction. Only terms with adjusted p-values lower than 0.05 are displayed. Terms are grouped based on their kappa score level (0.4) calculated using the ClueGO app and are represented by the most significant term in each group. Error bars represent ± SEM of biological replicates (n = 3). p: two-tailed Welch’s t-test. *p < 0.05, **p < 0.005, ***p < 0.0005. NS, not significant.

Interestingly, we noticed that many of the proteins encoded by the hit genes are among the top 1,000 proteins listed in Vesiclepedia^21,22^, a database compiling proteins frequently detected in/on sEVs (Fig. 3g, Supplementary Fig. 10), which implies that many proteins regulating sEV release are enriched in/on sEVs themselves. STRING analysis^23^ confirmed that the many of the hit genes, including those encoding the sEVs-resident proteins, were functionally connected to the validated genes as well as their neighbor genes and other hit genes previously reported as sEV release regulators, including VTI1B^24^, YKT6^25^ and ATP6V1A^26^ (Fig. 3h). Further, many of the hit genes also have biologically confirmed interactions with sEV release regulators such as PDCD6IP (ALIX)^27^, Rab5a^28^, Rab9a^28^, SMPD3^29^, ARF6^30^, and SNARE proteins^24^ (Supplementary Fig. 11). Together, these results demonstrate that genes regulating sEVs release are highly enriched in the CIBER screening hits.

We also performed gene ontology (GO) enrichment analysis^31,32^ of hit genes to extract putative biological processes relevant to sEVs release (Fig. 3i, Supplementary Data 5). It is noteworthy that the GO terms involved in vesicle-mediated transport are enriched in lower hits in this assay. Multiple previously unknown gene sets, such as those involved in tRNA metabolism (lower hits) and cholesterol metabolism (upper hits), are suggested to be involved in sEV release. Interestingly, GO terms grouped as “endosome organization”, including ESCRT proteins, were detected as strong upper hits in the screening, which we had not expected since they are known to be essential for sEV release^6,7^. However, validation with siRNA for two ESCRT members PTPN23 and VPS28 (Fig. 3f) indicated that depletion of ESCRT proteins could indeed upregulate sEV release in our setting, as reported in a different study^33^. The functions of ESCRT proteins are quite complex^34^ and some sEV release is known to be ESCRT-independent^35^, so we presume that ESCRT downregulation could act differently on sEV release depending on the experimental setting. In any case, these data support the conclusion that CIBER screening can comprehensively identify multiple genes and gene sets that significantly affect sEV release processes.

### Comparison of the regulators of CD63^+^ and CD9^+^ sEVs

sEVs are known to be heterogeneous, but how the release of different sEV subpopulations is controlled remains poorly understood^6^. We hypothesized that CIBER screening using different sEV markers to load gRNA would uncover the sEV release regulators in a subpopulation-specific manner.

Focusing on CD9 as another extensively used sEV marker, we conducted additional CIBER screening using CD9-dCas9 (CD9-CIBER hereafter; Supplementary Fig. 12, Supplementary Data 4). Firstly, we confirmed that CD9 was detected as a strong lower hit in the new screening, while CD63 was not (Fig. 4a), establishing the flexibility of our barcoding system to selectively load gRNA into the targeted sEV subpopulation. The z-REs of each gene in each screening are somewhat similar (R = 0.43 for all genes (Fig.4a), 0.63 for ESCRT genes (Supplementary Fig.13)) and about 30% of the hit genes overlapped (101 genes for lower hits and 64 genes for upper hits, Supplementary Fig. 14a), suggesting that there are many common regulators of CD63^+^ and CD9^+^ EV release. FASN and PI4KA were among the newly found shared lower/upper hits, and the effects of treatment with inhibitors of these proteins on CD63-nluc and CD9-nluc sEVs were confirmed to be similar (Fig. 3b,c, Supplementary Fig. 12c, d). It seems reasonable that multiple genes are common regulators of the release of CD63^+^ sEVs and CD9^+^ sEVs, considering that some proportion of sEVs should express both CD63 and CD9.

**Figure 4.**
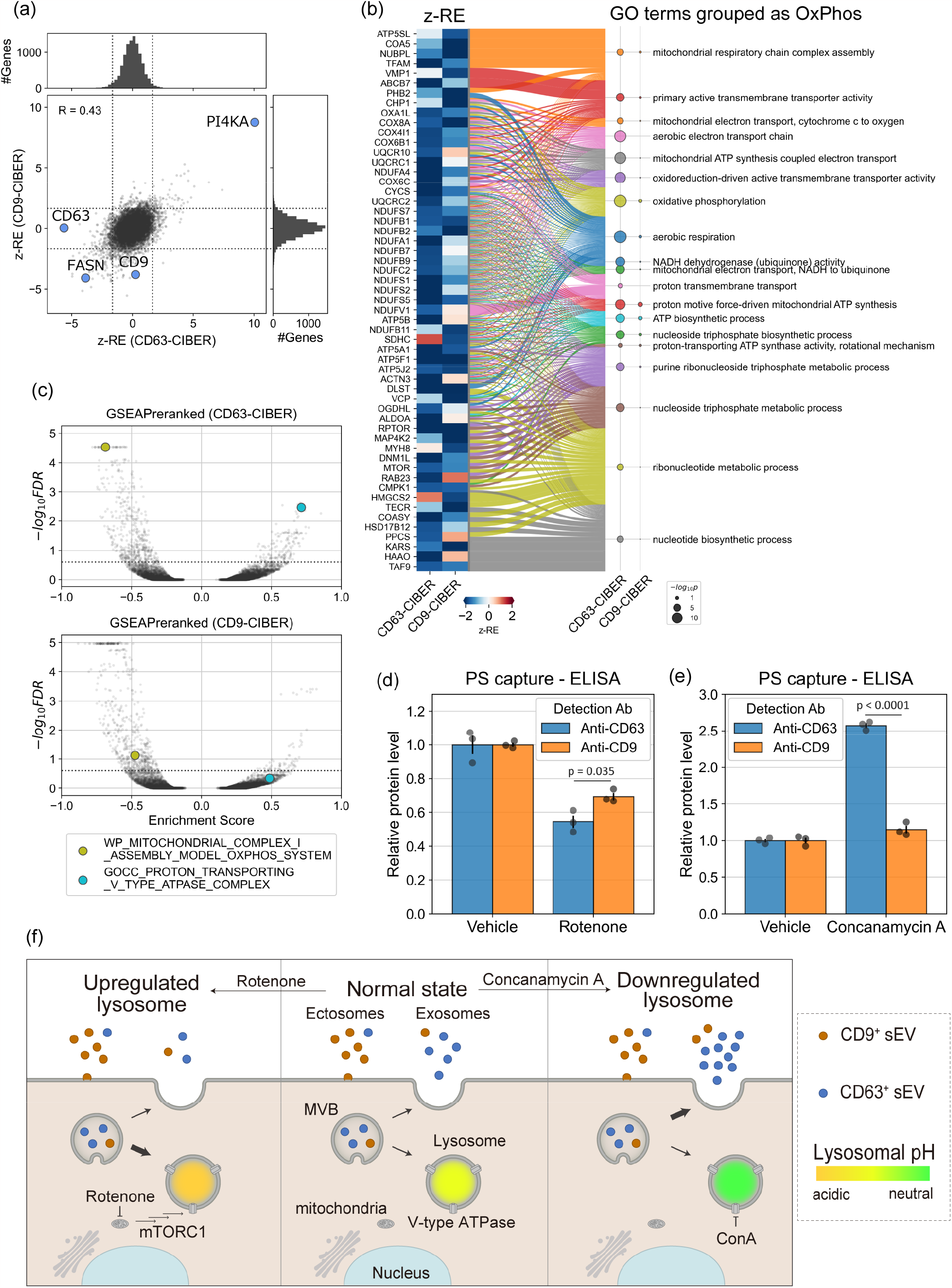
The results of the CD9-CIBER screening and comparison with those of CD63-CIBER screening. (a) The pair-wise plot of z-RE obtained from CD63/CD9-CIBER screen for 10,410 genes. The dashed lines show +/-1.65. (b) Gene-GO term association. The heatmap (left) shows z-RE of genes annotated to GO terms grouped as OxPhos in Fig. 3i. Only genes with z -RE less than -1.65 in either screening are displayed. The Sanky diagram (middle) indicates the connection of genes and their annotated terms. The scatter plot (right) shows the -log_10_(p-value) calculated for each GO term (Fisher’s exact test adjusted by Holm correction). (c) Volcano plot of enrichment score calculated via gene set enrichment analysis (GSEA). All the screened 10,410 genes were ranked by z-RE and analyzed using GSEAPreranked. Each plot represents a gene set. The dashed line shows the FDR (false discovery rate) of 0.25. In this data representation, KO or inhibition of components annotated to gene sets in the upper left/upper right area is predicted to downregulate/upregulate the release of each subpopulation of sEVs, respectively. More detailed results of the GSEA analyses are listed in Supplementary Data 6. (d) Relative abundance of CD63 and CD9 on the surface of sEVs after treatment with rotenone at 10 nM for 24 hours. sEVs released from wild-type HEK293T cells were captured on Tim4-immobilized wells and detected using either anti-CD63 antibody or anti-CD9 antibody conjugated with horseradish peroxidase (HRP). Vehicle: 0.1% DMSO. Error bars represent ± SEM of biological replicates (n = 3). p: two-tailed Welch’s t-test. (e) Relative abundance of CD63 and CD9 on sEVs surface after treatment with concanamycin A (ConA) at 1 nM for 24 hours, measured as shown in Fig. 5c. Error bars represent ± SEM of biological replicates (n = 3). p: two-tailed Welch’s t-test. (f) Proposed mechanisms of selective change of the release of CD63^+^ sEVs associated with lysosomal perturbation. Treatment of cells with the V-type ATPase inhibitor concanamycin A (ConA) neutralizes lysosomes, and the contents of MVB tend to evade lysosomal degradation, leading to increased release of “exosomal” CD63^+^ sEVs. Conversely, treatment of cells with an inhibitor of mitochondrial complex I, rotenone, enhances the activity of lysosome probably via mTORC1 and MIT/TFE signaling pathways, leading to decreased release of exosomal CD63^+^ sEVs. On the other hand, CD9^+^ sEVs are rather “ectosomal”, so their release is less affected by lysosomal perturbation.

We next set out to compare the results of CD63-CIBER and CD9-CIBER by GO enrichment analysis and gene set enrichment analysis (GSEA)^36^, aiming for the robust detection of factors that act differently on the release of different sEV subpopulations. Firstly, we noticed the appearance of significant GO terms grouped as oxidative phosphorylation (OxPhos) only in the lower hits of CD63-CIBER (Fig. 3i); only 2 out of 19 terms were significant for CD9-CIBER and genes annotated to these terms (mostly encoding mitochondrial proteins) were CD63-CIBER-specific lower hits (Fig. 4b, Supplementary Fig. 14b-e). We decided to pursue this further, because the relationship between sEV heterogeneity and mitochondrial ATP synthesis is not well understood. We conducted a parallel GSEA analysis by using z-REs of each screening as a pre-ranked query array, and the volcano plot of the analyzed ∼12,000 gene sets again suggested that inhibition of mitochondrial ATP biosynthesis would more significantly inhibit CD63^+^ sEV release (Fig. 4c). Indeed, the treatment of HEK293T cells with rotenone (a mitochondrial complex I inhibitor) strongly downregulated the release of CD63^+^ sEVs; this was confirmed with native sEVs by a sandwich ELISA of CD63/CD9 with phosphatidylserine on the sEV membrane (PS capture ELISA)^37^ (Fig. 4d). These results show that CIBER screening with different sEV markers can identify new sEV release regulators in a subpopulation-specific manner. On the other hand, the GSEA analysis suggested that inhibition of proton-transporting V-type ATPase would upregulate the release of CD63^+^ sEVs rather than CD9^+^ sEVs (Fig. 4c, Supplementary Data 6). Indeed, treatment of HEK293T cells with a potent V-type ATPase inhibitor, concanamycin A, markedly increased CD63^+^ sEV release (Fig. 4e). The result with concanamycin A is in line with the recent observation that CD63^+^ sEVs are “exosomal” (MVB-derived), while CD9^+^ sEVs are more “ectosomal” (plasma membrane-derived) in Hela cells, and V-type ATPase inhibition selectively downregulates degradation of sEVs produced in MVBs^38^. Conversely, we recently showed that inhibition of ATP biosynthesis in mitochondria could promote lysosomal function through mTORC1 and MIT/TFE signaling^39^. So, we believe our findings can be explained as follows: lysosomal function predominantly affects the release of more “exosomal” CD63^+^ sEVs, and thus the inhibition/upregulation of lysosomal function promotes/suppresses the release of CD63^+^ sEVs, respectively (Fig. 4f). This idea is consistent with the results of live-cell imaging showing that CD63 accumulates in intracellular vesicular compartments, while CD9 is mainly localized at the plasma membrane in HEK293T cells used in our study (Supplementary Fig. 15), and the fact that rotenone treatment activates cellular lysosomal function (Supplementary Fig.16).

As regards other new regulators that act differently on the release of CD63^+^ sEVs and CD9^+^ sEVs, the results of GSEA of CD63/CD9-CIBER and tf-idf (term frequency-inverse document frequency) analysis^40^ suggested that the downregulation of the cell cycle has a negative effect predominantly on the release of CD9^+^ sEVs (Fig. 5a, Supplementary Fig. 17; note that tf-idf analysis detected a stronger relationship of lysosomal activity with the release of CD63^+^ sEVs again). As expected, halting the cell cycle at the G2/M phase with a cyclin-dependent kinase inhibitor, dinaciclib, suppressed the release of only CD9^+^ sEVs (Fig. 5b, Supplementary Fig.18). In addition, we synchronized HEK293T cells expressing CD63-nluc or CD9-nluc in the S phase by imposing a double thymidine block (DTB) and periodically estimated the CD63^+^ and CD9^+^ sEVs in the culture supernatant by luminescence measurement while monitoring the progression of the cell cycle after the release from DTB (Fig. 5c, d). We found that the release of CD9^+^ sEVs, but not CD63^+^ sEVs, was negatively correlated with the ratio of cells at the M phase, and the release of CD9^+^ sEVs seemed to reach the maximum when most of the cells had completed cell division (Fig. 5e). These results are consistent with the idea that the release of CD9^+^ sEVs is synchronized with the cell cycle. To our knowledge, this study is the first to demonstrate an effect of the cell cycle on sEV heterogeneity.

**Figure 5.**
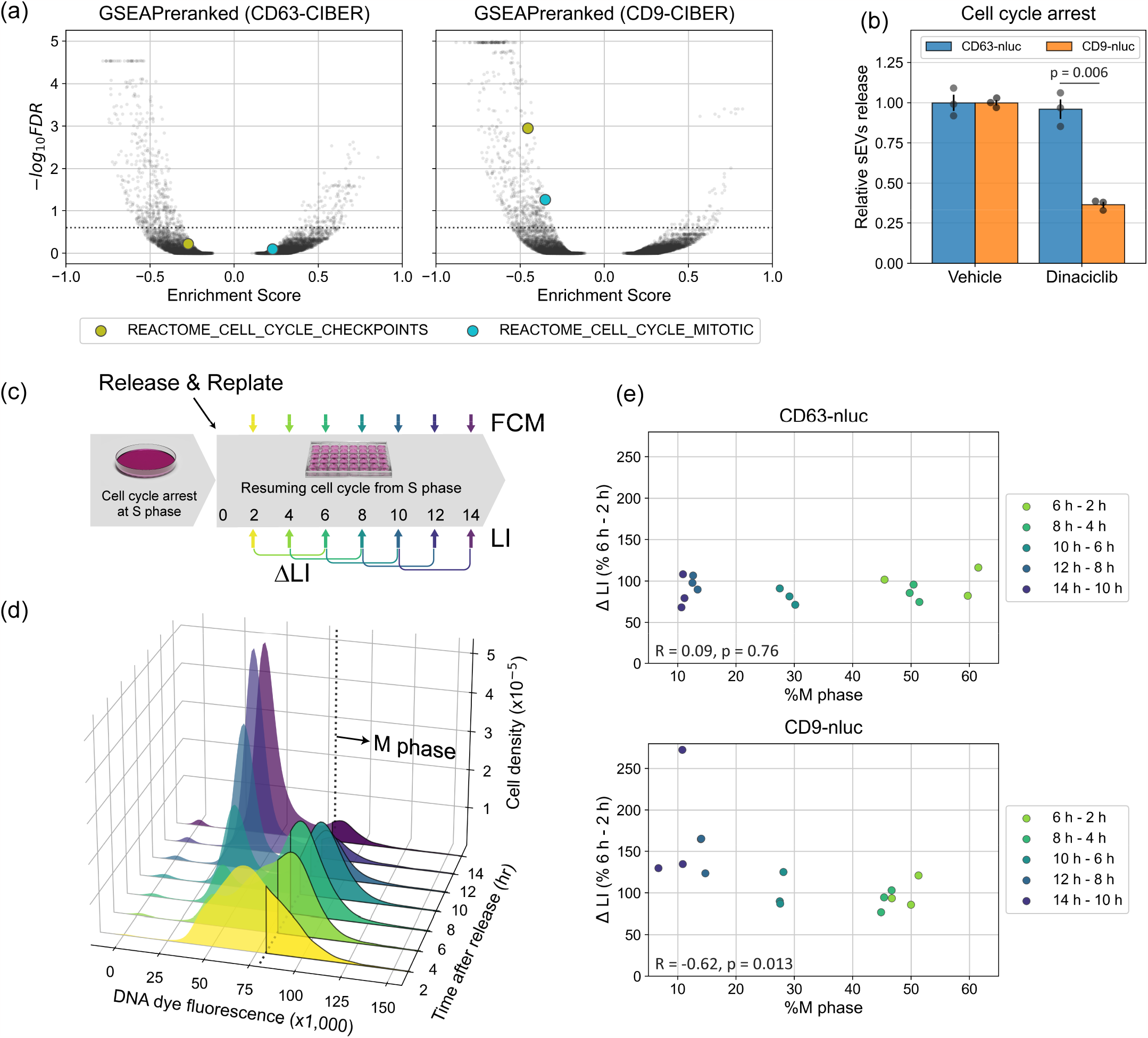
Synchronization of the cell cycle and release of CD9^+^ sEVs. (a) Volcano plot of enrichment score calculated via gene set enrichment analysis (GSEA). All the screened 10,410 genes were ranked by z-RE and analyzed using GSEAPreranked. Each plot represents a gene set. The dashed line shows FDR of 0.25. More detailed results of the GSEA analyses are listed in Supplementary Data 6. (b) Relative release of CD63^+^ or CD9^+^ sEVs during cell cycle arrest. HEK293T cells stably expressing CD63-nluc or CD9-nluc were treated with dinaciclib at 10 nM to stop the cell cycle at the G2/M phase for 24 hours. Cells were replated and additionally cultured in the presence of dinaciclib for 12 hours. LI of CM was measured at 4 and 12 hours after replating. Relative sEVs release was calculated as the difference of LI between the two time points. Error bars represent ± SEM of biological replicates (n = 3). p: two-tailed Welch’s t-test.(c) Time course of the cell-cycle synchronization assay. HEK293T cells expressing CD63-nluc or CD9-nluc are synchronized at S phase by means of a double thymidine block in bulk culture and then replated into separated wells without thymidine, allowing the cells to resume the cell cycle from the S phase. LI of CM and DNA abundance of the cells were measured at 2, 4, 6, 8, 10, 12, and 14 h after replacing the media. Three wells were used for each time point. (d) Representative result of flow cytometry (FCM) with CD63-nluc expressing cells shown in Fig. 5c. (e) Results of the synchronization assay. The percentage of cells in Fig. 5d with DNA dye fluorescence (x1,000) greater than 80 was defined as %M phase. ΔLI is the difference between the LI at a time point and the averaged value 4 hours earlier. p: t-test for correlation.

## Discussion

We have developed a high-throughput method named CIBER screening that enables the identification of sEV biogenesis/release regulators in a pooled manner and employed it to identify new genes and functionally linked gene clusters that affect sEV release. An especially noteworthy finding is that multiple pathways differentially affect the release of CD63^+^ and CD9^+^ sEVs.

To our knowledge, there has been only one previous study on pooled screening of sEV release regulators based on barcoded sEVs, which used miRNAs bearing short EV-targeting nucleotide tags as barcodes loaded in sEVs^9^. However, that study had significant limitations including the necessity for constructing complex custom-made libraries, a high sequence bias for sEV barcoding, the need for a huge amount of cell culture supernatant to harvest sEVs for analysis of the barcodes (>100 L for assaying the whole genome), and inability to examine the effect of cell viability and heterogeneity of sEVs. CIBER screening is unique in that CRISPR gRNAs are directly and actively loaded in sEVs as barcodes through interaction with dCas9 fused with an sEV marker. This feature enables efficient and uniform barcoding of sEVs with publicly available gRNA libraries, requiring only ∼400 mL of cell culture supernatant (to assay half the genome). In practice, this system allows a single experimenter to implement a large-scale screening within 2∼3 weeks with a ready-to-use gRNA-encoding lentivirus and stable cell lines expressing the necessary components (see Supplementary Fig. 19 for detailed comparison). The sEV-marker-driven gRNA loading into sEVs also allows subpopulation-specific analysis of sEV release for the first time. Furthermore, the ability of CIBER screening to cancel out the changes of barcode abundance in cells upon gene KO is very important, because analysis of gRNA abundance only in the sEV fraction in our screening failed to extract the factors differently affecting CD63 sEV release and CD9^+^ sEV release (Supplementary Fig. 20).

It should be noted that several proteins, including Rab family members (e.g. Rab11, Rab27a, Rab35), that reportedly regulate sEV biogenesis^41^ were not detected as hits by CIBER screening with the current settings. This could be due to the existence of genes having redundant functions or to inefficacy of the gRNAs used, but it is also possible that the KO of a single gene could induce compensation via other pathways, which would likely be influenced in part by the assay timeline. To address this issue, testing CIBER screening with CRISPR activation or inhibition should be an interesting option to explore other regulators, since the assay timeline with these systems is shorter than that with the CRISPR KO screen^42^.

Overall, we believe CIBER screening provides massive information on sEV release that would be inaccessible with conventional methods. Further, the present data should be a useful resource for future studies on sEVs, including more detailed analyses of how each gene regulates sEV release.

For the future, the different effects of the cell cycle on CD63^+^ and CD9^+^ sEV release raise a question about how cell-cycle-inhibiting anti-cancer drugs might influence the release of a specific subpopulation of sEVs that affects cancer malignancy. Pursuing this kind of question should deepen our understanding of the contribution of sEV heterogeneity to diseases. Related to this point, large-scale parallel CIBER screenings of cell-type-specific regulators of sEV would be an interesting approach to the discovery of novel drug targets for sEV-related diseases. Besides, the knowledge gained by CIBER screening could be applied for the more efficient production of sEV-based next-generation therapeutics with controlled heterogeneity. It might also be possible to apply CIBER screening to other classes of EVs including viruses, by changing the dCas9-recruiting protein. From a different viewpoint, the excellent correlation of gRNA compositions in the sEV and cellular fractions in our system implies that this sEV-barcoding system would also be applicable for continuously monitoring cell population dynamics with gRNA composition in sEVs without destroying the cells, allowing for continuous cell-free CRISPR screening (Supplementary Fig. 21; the CIBER screening can be used as a counter assay). Taking into consideration that sEVs protect the enclosed RNA, which reflects the state of the originating cells, from degradation *in vivo*^43,44^, the system could be potentially adapted even for continuous *in vivo* CRISPR screening. Thus, we envision that CIBER screening will be applicable for a variety of purposes in the future.

## Author contributions

K.K. and R.K. designed and conducted the majority of the experiments. K.K., R.K., and T.M. performed analysis and visualization of the screening data with a custom-made analysis pipeline. K.H., C.O., M.K., S.O., and Y.U. advised on the experiments. K.K. and R.K. co-wrote the manuscript with input from all the authors. R.K. conceived, directed, and supervised the entire project.

## Acknowledgment

We thank Prof. Kenzo Hirose, Dr. Shigeyuki Namiki and Dr. Daisuke Asanuma for providing the infrastructure necessary to conduct experiments using lentivirus, Dr. Kiyoshi Yamaguchi for the help for using Ion Proton, and the suppliers of the Addgene constructs used in this study. This work was supported by the Japan Science and Technology Agency (JST) PRESTO program (JPMJPR17H5 to R.K.), JST FOREST program (JPMJFR214N to R.K.), JST CREST program (JPMJCR19H1 to R.K. and S.O.), ATI research grant (to R.K.), and HFSP Career Development Award (CDA-00008/2019-C to R.K.). K.K. was supported by a Grant-in-Aid for JSPS fellows and WINGS-LST program of the University of Tokyo.

## Corresponding authors

Correspondence and requests for materials should be addressed to Ryosuke Kojima.

## Methods

### Cell culture

HEK293T cells were maintained in Dulbecco’s modified Eagle’s medium (DMEM, High Glucose with L-Glutamate, FUJIFILM Wako) with 10% (v/v) fetal bovine serum (FBS, Biosera) and 1 % (v/v) penicillin/streptomycin solution (PS, FUJIFILM Wako) at 37°C in a humidified atmosphere containing 5% CO_2_.

### Transfection

Cells were plated at 2.5 × 10^5^ cells/mL onto a 10 cm dish in 10 mL medium and cultured for 24 hours before transfection. Ten µg of total DNA in 1 mL of plain optiMEM was mixed with 40 µl of Polyethyleneimine “Max” (PEI, Polyscience #24765, 1 mg/mL in dH_2_O), briefly vortexed and incubated at room temperature for 15 min. Cell culture medium was renewed before transfection and DNA/PEI mixture was added dropwise to the culture. After a sufficient cultivation period (typically 8-16 h), the medium was renewed again for downstream applications. Transfections were scaled up or down based on the culture area (cm^2^) when necessary.

### Establishment and maintenance of stable cell lines

The Sleeping Beauty transposase system^45,46^ was mainly used for establishing stably transfected cell lines, except for the lentivirus infection of gRNA cassettes. Cells were transfected with DNA mix containing 50 ng of transposase-encoding plasmid (addgene #34879, pCMV(CAT)T7-SB100) and 950 ng of transposon plasmid on a 12-well plate and selected in antibiotics-containing medium (10 µg/mL blasticidin and 300 µg/mL hygromycin). Sufficiently expanded cells were further sorted by FACS (FACS Aria II or III, BD) and the top 10% of cells with strong fluorescence were collected. Established cell lines were maintained in the selecting medium.

### Plasmid construction

The cloning strategy and oligo DNA used for plasmid construction are described in Supplementary Tables 1 and 2.

### Lentivirus production, titration, and transduction to cells

Production and titration of lentivirus were conducted as previously reported^42^ with some modifications. In brief, Lenti-X cells (Clontech) were transfected with pMD2.G (addgene #12259), psPAX2 (addgene #12260) and the pooled gRNA library at the ratio of 1:2:4 (ng) and cultured for 6 hours. Then, the medium was renewed and culture was continued for 42 hours. Virus-containing medium was passed through a 0.45 µm filter (cat. #SLHVR33RB, Millipore), aliquoted and stored at -80°C. We performed lentivirus production on a well of a 12-well plate for a single gRNA and on a 500 cm^2^ dish for DTKP, PROT, ACOC, TMMO libraries^16^ (spacer sequences are listed in Supplementary Data 7). For titration, cells mixed with polybrene (cat. #12996-81, nacalai tesque, 8 µg/mL final concentration) were transduced by spinfection (1000 x g for 2 hours at 33 °C) with increasing volumes of lentivirus and then selected with 0.3 µg/mL puromycin for 2-3 days under normal culture conditions. The multiplicity of infection was determined by comparing the viability of selected cells with that of the no-puromycin control using a Cell Counting Kit-8 (cat. #CK04, Dojindo).

For screening, HEK293T cells were transduced at the MOI of 0.3 according to the same protocol. The number of cells to be transduced was calculated according to the following equation to ensure that at least 500 cells were transduced with each gRNA in a library: (the number of gRNA in library) x 500 / 0.3. After the transduction, cells were selected for 7 days to achieve maximal knockout and used for sEVs isolation while maintaining 500 cells/gRNA.

### Isolation of sEVs

Cells plated at 1.5-2.0 × 10^5^ cells/mL were propagated to 70-80% confluency for sEVs production. The medium was changed to optiMEM with 1% (v/v) P/S and cells were cultured for another 24-36 hours. The culture supernatant was collected and centrifuged stepwise at 300 x g for 5 min and 2,000 x g for 10 min to remove cells and large debris, followed by filtration through a 0.22 µm filter (cat. #SLGV033RS Millipore) to remove small debris. At this time, cells were washed with PBS and lysed in TRIzol (cat. #15596018, Invitrogen) at 10^7^ cells/mL after collection of the culture supernatant for screening. Filtered supernatant was transferred to tubes for ultracentrifugation (cat. #344058, Beckman Coulter) to which 200 µL of Optiprep (Abbott Diagnostics Technologies AS) was added at the bottom as a cushion^47^. Small EVs were isolated by ultracentrifugation at 120,000 x g and 4°C for 120 min using Optima XE-90 equipment with a SW32Ti rotor (Beckman Coulter). Supernatants were discarded using a pipette, leaving 1.5-2.0 mL of the sEVs fraction at the bottom. The pellets were resuspended in vesicle-depleted PBS and centrifuged again with 50 µL of Optiprep cushion. Supernatants were discarded, leaving ∼0.5 mL of the sEVs fraction. The resulting pellets were resuspended in vesicle-depleted PBS and transferred into PROKEEP Protein Low Binding Tubes (cat. #PK-15C-500N, Watson). We usually resuspended sEVs from 100 mL of culture supernatant in 1 mL of PBS, typically resulting in 5-1.0 × 10^11^ particles/mL. Washed sEVs were kept at 4°C for short-term storage (up to a week) or flash-frozen and kept at -80°C for longer-term storage.

### Nanoparticle tracking analysis (NTA)

Nanosight LM10 equipment (Malvern Panalytical) was used for NTA followed by evaluation using the NTA software (ver. 3.2). The analysis was performed using a 488 nm laser with a recording time of 30 seconds, camera level of 15-16 and detection threshold of 5. Three recordings were sequentially performed for each sample. Samples were diluted to 10^9^ ∼ 10^10^ particles/mL before recording. When measuring culture supernatant, samples were cleared by centrifugation at 300 x g for 5 minutes, 2,000 x g for 10 minutes and 10,000 x g for 30 minutes before the measurement.

### Evaluation of gRNA amount in sEVs

Wild-type HEK293T cells or HEK293T cells stably expressing CD63-dCas9 were transduced with lentivirus carrying gRNA. After selection, sEVs were isolated from 30 mL of culture supernatant as described above. The concentration of sEVs were measured by NTA. Next, samples were diluted to 1 × 10^11^ particles/mL. Three µL aliquots of diluted samples were lysed by mixing with 3 µL of 0.1% Triton X-100 in Nuclease-Free Water (NFW, cat. #AM9937, Invitrogen) and incubating for 10 minutes at room temperature. The Ct values of samples were determined by quantitative PCR (qPCR) using a Luna Universal Probe One-Step RT-qPCR Kit (cat. #E3006, New England Biolabs) with LightCycler 480 (Roche Diagnostics) according to the manufacturers’ instructions. The sequences of primer and probe (IDT PrimeTime Mini qPCR assay) are listed on Supplementary Table 2. The same reaction was performed with each gRNA-containing plasmid (in pCRISPRia-v2) at 0.0001 – 1000 pg/µL to obtain a standard plot which was used to calculate the absolute number of gRNA molecules in each sample. The number of gRNA molecules was divided by the number of particles in the PCR reaction to calculate copies/particle (Fig. 2c).

### Sample preparation for NGS

#### -RNA extraction and DNA digestion

For cellular RNA, total RNA was extracted from at least 500 cells/gRNA according to the manufacturer’s protocol and dissolved in 40 µL of NFW. Contaminant DNA in 50 µg of extracted RNA was digested with DNaseI (cat. #2270A, Takara) according to the manufacturer’s protocol. After DNA digestion, the reaction was quenched by adding 5 µL of EDTA-2Na [250 mM] followed by incubation at 80°C for 2 minutes. Then 45 µL of NFW, 10 µL of NaOAc [3 M] and 250 µL of chilled EtOH were added to precipitate nucleic acids, and the mixture was incubated at -80°C for 20 minutes, and centrifuged at 120,000 x g and 4°C for 30 min. The resulting pellet was washed with 750 µL of chilled 70% EtOH and resuspended in 40 µL of NFA. Two µg of RNA was used for reverse transcription (RT).

For sEVs, total RNA was extracted from 250 µL of sEVs solution with 1 mL of TRIzol. The RNA-containing phase was mixed with GlycoBlue™ Coprecipitant (cat. #AM9515, Invitrogen) before isopropanol precipitation. The RNA pellet was dissolved in 5 µl of NFW and used for RT.

#### -RT of transcribed gRNA (SMART technology^48^) and clean-up

Five µL of RNA was mixed with 0.5 µL of oKK145 [12 µM], incubated at 70°C for 3 min, put on ice to form RNA/primer complex and mixed with 4.5 µL of RT solution {0.5 µL of SMARTScribe RTase (cat. #639536, Takara), 2 µL of 5x first strand buffer (supplied with RTase), 0.25 µL of DTT (supplied with RTase), 1 µL of dNTPs (cat. #N0447, NEB), 0.5 µL of custom LNA-TSO [12 µM] (cat. # 339412, Qiagen) and 0.25 µl of RNase inhibitor (cat. #2313A, Takara)}. The RT mixture was incubated at 42°C for 120 min and heated at 70°C for 10 min. After the RT reaction, 18 µL of AMPure XP (cat. #BC-A63881, Beckman Coulter) was added to the solution and incubated for 10 min at room temperature. The cDNA bound to beads was separated on magnet rack and subjected to tagging PCR without washing. For MS2-gRNA, Oligo #4 was used instead of oKK145.

#### -Tagging PCR

To the bead-bound DNA, 25 µL of PCR solution {0.15 µL of oKK147 [50 µM], 0.15 µL of (each of Oligo #6-Oligo #17) [50 µM], 12.5 µL of Tks Gflex DNA polymerase Low DNA (cat. #R091A, Takara) and 12.2 µL of NFW} was added. PCR conditions were as follows: 95°C for 1 minute, 20 cycles of (98°C for 10 seconds, 65°C for 30 seconds, 68°C for 30 seconds), 68°C for 30 seconds and 4°C hold. The PCR product (NGS sample) was purified with 35 µL of AMPure XP and eluted in 20 µL NFW. The concentration of NGS sample was measured by qPCR using a primer set of oKK120/oKK121. The standard curve was obtained using a commercial control (*E. coli* DH10B library control in Ion Library TaqMan™ Quantitation Kit, Thermo Fisher Scientific Inc.) diluted to 6.8, 0.68, 0.068, 0.0068, and 0.00068 pM. For MS2-gRNA, Oligo #5 was used instead of oKK147.

### Electrophoresis on the Agilent 2100 Bioanalyzer

The NGS sample was diluted with TE buffer and assayed using a High Sensitivity DNA Kit (cat. #5067-4626, Agilent Technologies) on a Bioanalyzer 2100. Raw data was exported as a csv file using the 2100 expert software.

### NGS using Ion Proton

Each sample was diluted to 50 pM and sequenced by an Ion Proton instrument (Thermo Fisher Scientific Inc.) using an Ion PI Hi-Q Chef Kit (cat. #A27198) and Ion PI Chip Kit v3 (cat. #A26771). One chip generally offers 10^8^ valid reads, so we pooled 4 samples (25,000-30,000 gRNAs/sample) aiming for >500 reads/gRNA for each sequencing. The chip was prepared using an Ion Chef instrument (Thermo Fisher Scientific Inc.). FileExporter was used to export fastq files for downstream analysis.

### NGS Data processing

We share a Python script count_barcodes.py (a modified version of count_spacers.py^42^) for barcode counting (see the section of Code availability). We prepared a .xlsx file containing the barcode id in each line to be referenced with a header named ‘id’. The format of the barcode id should be ‘{target gene}_{spacer sequence}_{additional information}’ (e.g., ‘AADACL2_GAAAGTCAGAAACCCGA_2832.7_DTKP’). Each read is assigned to a corresponding barcode if the 8 bp of scaffold sequence flanked to the spacer (GTTTAAGA; substituted to KEY in the script) is detected and 17 bp of the sequence upstream of KEY is identical to the spacer sequence. All the generated count files are available in Supplementary Data 2. Relevant statistics are also available in Supplementary Data 1.

### Calculation of z-normalized release effect (z-RE)

We share a Python script calculate_zRE.py for calculation of z-normalized release effect (see the section of Code availability). Count files on Supplementary Data 2 are compatible with calculate_zRE.py. Raw read counts are normalized to calculate nRC so that average nRC in a sample is 1. Each barcode would have 4 nRCs depending on the sample origin; (cell/sEVs) x (Cas9+/Cas9-). Barcodes with nRC lower than 0.05 in at least one sample were excluded from downstream calculation and after this exclusion, any genes with less than 3 barcodes were also excluded. FC_cells_ and FC_sEVs_ are calculated for each barcode as log_2_(nRC_cell x Cas9+_/nRC_cell x Cas9-_) and log_2_(nRC_sEVs x Cas9+_/nRC_sEVs x Cas9-_), respectively. Linear regression was performed on the scatter plot with FC_cells_ on the x-axis and FC_sEVs_ on y-axis. The release effects (RE) for each barcode were tentatively calculated as residues from the regression line (RE at gRNA level). Among every barcode group targeting the same gene, the barcodes with highest/lowest RE were excluded from the scatter plot and the regression line was drawn again. RE at the gRNA level was calculated again using the new regression line. RE at the gene level is the median value of REs of barcodes targeting the gene. REs at the gene level were z-normalized for each subpool library to calculate z-RE. Genes with z-RE larger than 1.65 are regarded as upper hits and genes with z-RE lower than -1.65 are regarded as lower hits in this paper.

### Inhibition assay with GSK-A1 and C75

Cells were plated onto 48-well plates (4.5 × 10^4^ cells in 300 µL of supplemented media) and allowed to expand to around 70% confluency. Then, media were replaced with optiMEM (1% PS) containing GSK-A1 (cat. #SYN-1219-M001, AdipoGen) or C75 (cat. #10005270, Cayman Chemical Co.). Cells were treated for 24 hours for GSK-A1 or for 30 minutes followed by transfer to supplemented media without inhibitor for C75. Wild-type HEK293T cells were used for nanoparticle tracking analysis and HEK293T cells expressing CD63-nluc or CD9-nluc were used for nluc-based reporter assay.

For reporter assay, culture supernatant was centrifuged at 300 x g for 5 min and 2,000 x g for 10 minutes to remove cell debris. The resulting supernatant was diluted at 1:50 with PBS and used for luminescence measurement with the Nano-Glo Luciferase Assay System (cat. #N1110, Promega) according to the manufacturer’s protocol. The protein concentration of the cellular fraction was used to normalize the measured luminescence intensity (LI). After the removal of the culture supernatant, 75 µL of CelLytic M (cat. #C2978, Sigma Aldrich) was added to each well. The plate was gently agitated for 20 minutes at room temperature and the protein concentration was measured using a Pierce BCA Protein Assay Kit (cat. #0023227, Invitrogen) according to the manufacturer’s protocol. The LI was divided by the protein concentration of corresponding well to calculate normalized LI.

### Hit validation with siRNA

A reverse transfection protocol was adopted for siRNA transfection. The transfection mix was prepared by mixing 2 µL of Lipofectamine RNAiMAX Transfection Reagent (cat. #13778030, Thermo Fisher Scientific) in 98 µL of plain optiMEM and 20 µL of siRNA [500 nM] in 80 µL of plain optiMEM. Two hundred µL of transfection mix was incubated for 5 minutes at room temperature, and mixed with 800 µL of cell suspension at 2.0 × 10^5^ cells/mL and the cells were plated onto a well of a 12-well plate. Media were refreshed after 24 hours. Culture was continued for another 24 hours, and the cells were passaged to 3 wells of a 48-well plate and cultured for 24 hours. Media were replaced with optiMEM (1% PS) and incubation was continued for 24 hours. After the final incubation, the culture supernatant was harvested and processed as described above. The source information of siRNA is listed in Supplementary Table 4. All the siRNAs were used as a mixture of siRNA #1-siRNA #3. The 27-mer synthetic double-strand RNAs (DsiRNA, manufactured by IDT) were used for knocking down OSBP, TMED10 and GOLGA2 while conventional 21-mer siRNAs were used for other genes.

### Protein-protein interaction

Upper hits and lower hits were separately queried in the STRING database via the StringApp plugin (https://apps.cytoscape.org/apps/stringapp) on Cytoscape^49^ (ver. 3.9.1, https://cytoscape.org/). Interaction maps were drawn with default settings. Gene products were represented as nodes and connected with edges to each other if STRING analysis predicted interactions. Nodes without any interactions were not displayed. Node filling colors were changed to show z-RE. The borders of the nodes for genes validated in Fig. 3f (OSBP, FASN, TMED10, GOLGA2, PTPN23, CAB39, VPS28, PI4KA) were changed to thick yellow. The borders of the nodes directly connected to validated genes were changed to thick black.

### Gene ontology (GO) Analysis

Enrichment analysis against the GO Biological Process data set was performed using the ClueGO plugin (ver. 2.5.9, https://apps.cytoscape.org/apps/cluego) on Cytoscape (ver. 3.9.1). Analysis parameters were as follows; Marker Lists: upper hits or lower hits, Ontology:

GO_BiologicalProcess-EBI-UniProt-GOA-ACAP-ARAP_25.05.2022_00h00, p-value cutoff = 0.05, Correction Method = Bonferroni step down, Min GO Level = 6, Max GO Level = 13, Number of Genes = 2, Min Percentage = 5.0, GO Fusion = true, GO Group = true, Kappa Score Threshold = 0.4. We used raw output file (Suppl Data 5_GO analysis.xlsx, sheet ‘CD63low’, ‘CD63up’, ‘CD9low’ and ‘CD9up’) to draw Fig. 3i and S13. For Fig. 4b, OxPhos terms were individually analyzed with CD9-CIBER hits as query. The output results were added to the raw output file on sheet ‘CD9low’ and p-value adjustment by Bonferroni step down was manually performed to calculate adjusted p-value.

### Hit validation with PS (phosphatidylserine) capture ELISA

HEK293T cells were plated onto 48-well plates (4.5 × 10^4^ cells in 300 µL of supplemented media). Cells were allowed to expand to around 70% confluency. Then, media were replaced with 500 µL of supplemented media containing rotenone (cat. #R0090, Tokyo Chemical Industry) at 10 nM or concanamycin A (cat. #BVT-0237-C025, AdipoGen) at 1 nM. Cells were treated for 24 hours, then the culture media were harvested and cleared by stepwise centrifugation of 300 x g for 5 minutes, 2,000 x g for 10 minutes and 10,000 x g for 30 minutes at 4°C.

The expression levels of CD63 and CD9 on the sEVs surface were measured using a PS Capture Exosome ELISA Kit (cat. #298-80601, Fujifilm) according to the manufacturer’s protocol. Samples were diluted 1:10 with Reaction Buffer. Anti-CD63 antibody was supplied with the kit. Anti-CD9 antibody was purchased from Fujifilm (cat. # 014-27763) and used at 240 ng/mL diluted with Reaction Buffer.

### Gene set enrichment analysis (GSEA)

GSEA was performed with GSEA software (ver. 4.2.3, https://www.gsea-msigdb.org/gsea/index.jsp) using the following .gmt files (gene sets); h.all.v2023.1.Hs.symbols.gmt [Hallmarks], c2.all.v2023.1.Hs.symbols.gmt [Curated], c5.all.v2023.1.Hs.symbols.gmt [Gene ontology]. Gene names and corresponding z-REs were queried to run GSEAPreranked with the default setting of 1000 permutations and No_Collapse.

### Cell cycle arrest with dinaciclib

HEK293T cells expressing CD63-nluc or CD9-nluc were plated onto a 10 cm dish (1.5 × 10^6^ cells in 10 mL of supplemented media) and allowed to expand to around 70% confluency, then treated with dinaciclib (cat. # D479725, Toronto Research Chemicals) at 10 nM or vehicle (0.1% DMSO) in supplemented media for 24 hours. Cells were detached from the dish, diluted to 1.0 × 10^6^ cells/mL with supplemented media, and replated (500 µL) onto 6 wells of a 48-well plate. The cells were kept in dinaciclib media or vehicle until replating was finished. After 4 and 12 hours, the culture supernatant was harvested from 3 wells for each time point and used for LI measurement as described above. Relative EV release was calculated by subtracting the LI at 4 hours from the LI at 12 hours and normalizing the value to the mean value under vehicle conditions (Fig. 5b).

Immediately after media collection, cells were washed with 100 µL of PBS, resuspended in 250 µL of PBS, mixed with 2.5 µL of Cell Cycle Assay Solution Blue (cat. #C549, Dojinbo) and incubated for 15 minutes at 37°C in the dark to stain nuclei. Stained cells were directly subjected to flow cytometry (FCM) analysis on a BD LSR II (Becton, Dickinson and Company) to measure the DNA amount in individual cells. FCM data was processed on FACS Diva software (ver. 4.1).

### Double thymidine block

HEK293T cells were synchonized at the G1/S boundary by the double thymidine block (DTB) method as previously described^50^. DTB was performed by adding thymidine to a final concentration of 2 mM to a culture of 2.0 × 10^6^ HEK293T cells expressing CD63-nluc or CD9-nluc plated onto a 10 cm dish 24 hours prior to thymidine addition, incubating each dish for 18 hours, refreshing the media after washing the dish 3 times with PBS and incubating for 9 hours, and treating cells again with thymidine at 2 mM for 18 hours to complete cellular synchronization.

After the DTB, cells were detached from the dish, resuspended in supplemented media without thymidine, diluted to 1.0 × 10^6^ cells/mL, and replated (500 µL) onto 21 wells of a 48-well plate. At 2, 4, 6, 8, 10, 12 and 14 hours after replating, the LI of culture media and DNA amount in cells were measured in the same way as described for dinaciclib assay (n = 3). ΔLI is the relative sEV amount released during the 4-hour interval, calculated by subtracting the mean LI at 2, 4, 6, 8, 10 hours from LI of every replicate at 6, 8, 10, 12, 14 hours, respectively, and normalizing all values to mean ΔLI between 6 hours and 2 hours. FCM data processed on FACS Diva software were further analyzed using a Python package FlowCal^51^ (https://taborlab.github.io/FlowCal/) to calculate the ratio of cells at M phase. The G1 peak appeared around 45,000 < DNA dye fluorescence < 50,000 and the G2/M peak appeared around 85,000 < DNA dye fluorescence < 90,000, so we assigned cells with DNA dye fluorescence larger than 80,000 to M phase.

### Statistical analysis

Unless otherwise stated, all computational and statistical analyses in this study were performed using Excel, Python or R. Statistical details for each experiment can be found in the figure legends or corresponding method section. Differences with a p-value < 0.05 were considered significant.

## Code availability

We share python scripts used in this study via GitHub: https://github.com/Ryosuke-Kojima/CIBER-screening-paper

