## Supplementary Tables 1-4, Supplementary Figures 1-21 for "Barcoding of small extracellular vesicles with CRISPR-gRNA enables comprehensive, subpopulation-specific analysis of their biogenesis/release regulators"

### Inventory

**Supplementary Table 1.** Plasmids used in this study……………………………………………………………….3
**Supplementary Table 2.** Oligonucleotides used in this study…………………………………………………………4
**Supplementary Table 3.** Barcode information of NGS primers……………………..……………………………...6
**Supplementary Table 4.** Sources of siRNA used in this study…………………………...……………………7
**Supplementary Fig. 1.** Comparison of pooled CRISPR screening and CIBER screening……………………….…….8
**Supplementary Fig. 2.** Description and performance of MS2-based gRNA loading system…………..……………9
**Supplementary Fig. 3.** Results of nanoparticle tracking analysis (NTA) for sEVs isolated from culture media (CM) of HEK293T cells expressing each component…………………………………………………………………….9
**Supplementary Fig. 4.** Confirmation that co-expression of Cas9 has little effect on coverage of gRNAs in sEVs.10
**Supplementary Fig. 5.** Confirmation that CD63-dCas9 expression has little effect on Cas9-induced gene KO…..10
**Supplementary Fig. 6.** gRNA libraries used in this study………………………………………………………11
**Supplementary Fig. 7.** Comparison of fold-change of barcodes in cellular fraction and sEVs fraction…………...12
**Supplementary Fig. 8.** Concentration of sEVs released from HEK293T cells treated with GSK-A1 or C75……..12
**Supplementary Fig. 9.** Confirmation of gene knockdown by siRNA used for validation in Figure 3f……………13
**Supplementary Fig. 10.** Matching of the hit genes of CD63-CIBER and top 1,000 sEVs-associated proteins compiled from VesiclePedia database……………………………………………………………………………14
**Supplementary Fig. 11.** Connection of the hit genes in CD63-CIBER with known sEV release regulators….15
**Supplementary Fig. 12.** Results of CD9-CIBER screening……………………………………………………..16
**Supplementary Fig. 13.** z-RE of ESCRT genes obtained from CD63-CIBER and CD9-CIBER…………………17
**Supplementary Fig. 14.** Detailed analysis of OxPhos terms and their annotated genes…………………………...18
**Supplementary Fig. 15.** Subcellular localization study of CD63 and CD9…………………………..…………..19
**Supplementary Fig. 16.** Confirmation of lysosomal activity upon treatment with inhibitors……………………...20
**Supplementary Fig. 17.** Term frequency - inverse document frequency (TF-IDF) analysis of hit terms…………21
**Supplementary Fig. 18.** Flow cytometry analysis of cell cycle arrest by dinaciclib………………………………22
**Supplementary Fig. 19.** Comparison of sequencing results of CIBER screening and a previous study…………...23
**Supplementary Fig. 20.** Volcano plot of enrichment score calculated via GSEAPreranked with z-$FC_{sEVs}$………24
**Supplementary Fig. 21.** Data showing the applicability of the sEV-barcoding system for cell-free CRISPR screening………………………………………………………………………………………………………..25

**Supplementary methods**……………………………………………………………………………….…..26

**Supplementary references**……………………………………………………………………...…….28

| Plasmid | Description and Cloning Strategy | Reference /Source |
|---|---|---|
| **pKK47** | Constitutive expression of CD63-MS2. CD63-MS2 was PCR-amplified from pSA_Exo010 (unpublished, encoding MS2 reported by Wroblewska et al[1]) from using oKK29 and oKK103, digested with SfiI and cloned into the corresponding sites (SfiI) of pSBbi-GH (addgene #60514[2]). | This work |
| **pKK49** | Constitutive expression of mock MS2-gRNA for lentiviral transduction. gRNA scaffold bearing MS2 coat protein-binding motif was PCR-amplified from sgRNA(MS2) cloning backbone (addgene #61424[3]) using oKK109 and oKK105, digested with BlpI/XhoI and cloned into the corresponding sites (BlpI/XhoI) of pCRISPRia-v2 (addgene #84832[4]). | This work |
| **pKK59** | Constitutive expression of CD63 with EFGGGGSGTIS linker at the C-terminal. CD63 was PCR-amplified from pDB30[5] using oKK127 and oKK128, digested with SfiI and cloned into the corresponding sites (SfiI) of pSBbi-GH (addgene #60514[2]). The GTIS sequence in the linker can be digested by KpnI-HF/NheI-HF to allow insertion of the gene to be fused to CD63 | This work |
| **pKK60** | Constitutive expression of CD63-dCas9. dCas9 was PCR-amplified from lenti-EF1α-dCas9-KRAB-Puro (addgene #99372[6]) using oKK129 and oKK130 (excluding N-terminal nuclear localization signal (NLS), KRPAATKKAGQAKKKK), digested with KpnI-HF/NheI-HF and cloned into the corresponding sites (KpnI-HF/NheI-HF) of pKK59. | This work |
| **pRK300** | Constitutive expression of Cas9. Cas9 was PCR-amplified from lentiCas9-Blast (addgene #52962[7]) using oRyo004 and oRyo005, digested with SfiI and cloned into the corresponding sites (SfiI) of pSBbi-RB (addgene #60522[2]). | This work |
| **pKK89** | Constitutive expression of non-targeting gRNA (spacer sequence; GAAAGAGGGTCCCCCTGTAG). The non-targeting spacer was inserted by annealing oKK206 and oKK207 and cloned into pCRISPRia-v2 (addgene #84832[4]) which was digested with BlpI/BstXI to remove dummy spacer. The spacer sequence was selected from the Bassik human deletion library[8]. | This work |
| **pKK90** | Constitutive expression of non-targeting gRNA (gRNA #1) (spacer sequence; GAACAAGTTCCAATTTGTAT).oKK208 and oKK209 were inserted to pCRISPRia-v2 (addgene #84832[4]) in the same way as pKK89. The spacer sequence was selected from the Bassik human deletion library[8]. | This work |
| **pKK209** | Constitutive expression of BFP-targeting gRNA (gRNA #2) (spacer sequence; GCACATGAAGCTGTACATGG). oKK531 and oKK532 were inserted to pCRISPRia-v2 (addgene #84832[4]) in the same way as pKK89. The spacer sequence was designed using CHOPCHOP[9]. | This work |
| **pKK210** | Constitutive expression of BFP-targeting gRNA (spacer sequence; GGCGAAGGCAAGCCCTACGA). oKK533 and oKK534 were inserted to pCRISPRia-v2 (addgene #84832[4]) in the same way as pKK89. The spacer sequence was designed using CHOPCHOP[9]. | This work |
| **pKK104** | Constitutive expression of CD9 with EFGGGGSGTIS linker at the C-terminal. CD9 was PCR-amplified from pDB96[5] ($P_{hCMV}$-CD9-nluc-$pA_{bGH}$) using oKK620 and oKK621, digested with SfiI and cloned into the corresponding sites (SfiI) of pSBbi-GH (addgene #60514[4]). The GTIS sequence in the linker can be digested by KpnI-HF/NheI-HF to allow insertion of the gene to be fused to CD9. | This work |
| **pKK106** | Constitutive expression of CD9-dCas9. dCas9-encoding DNA fragment was collected from pKK60 digested with KpnI-HF/NheI-HF followed by agarose gel purification and inserted into the corresponding sites (KpnI-HF/NheI-HF) of pKK104. | This work |
| **pKK108** | Constitutive expression of CD63-nluc. nluc was PCR-amplified from pDB30[5] ($P_{hCMV}$-CD63-nluc-$pA_{bGH}$) using oKK236 and oKK237, digested with KpnI-HF/NheI-HF and cloned into the corresponding sites (KpnI-HF/NheI-HF) of pKK59. | This work |
| **pKK147** | Constitutive expression of CD9-nluc. nluc-encoding DNA fragment was collected from pKK108 digested with KpnI-HF/NheI-HF and inserted into the corresponding sites (KpnI-HF/NheI-HF) of pKK104. | This work |
| **pKK148** | Constitutive expression of CD63 with EFGGGGSGTIS linker at C-terminal. CD63-linker-encoding DNA fragment was collected from pKK59 digested with SfiI followed by agarose gel purification and inserted into the corresponding sites (SfiI) of pSBbi-Hyg (addgene #60524[4]). | This work |

| | | |
|---|---|---|
| **pKK149** | Constitutive expression of CD9 with EFGGGGSGTIS linker at the C-terminal. CD9-linker-encoding DNA fragment was collected from pKK104 digested with SfiI followed by agarose gel purification and inserted into the corresponding sites (SfiI) of pSBbi-Hyg (addgene #60524[4]). | This work |
| **pRK397** | Constitutive expression of CD63-mScarlet. mScarlet was PCR-amplified from pmScarlet_C1 (addgene #85042[10]) using oRK164 and oRK165, digested with KpnI-HF/NheI-HF and cloned into the corresponding sites (KpnI-HF/NheI-HF) of pKK148. | This work |
| **pKK151** | Constitutive expression of CD9-sfGFP. sfGFP was PCR-amplified from pHRdSV40-scFv-GCN4-sfGFP-VP64-GB1-NLS (addgene #60904[11]) using oKK334 and oKK335, digested with KpnI-HF/NheI-HF and cloned into the corresponding sites (KpnI-HF/NheI-HF) of pKK149. | This work |

**Supplementary Table 1.** Plasmids used in this study.

| **Name** | **Usage** | **Sequence (5' -> 3')** |
|---|---|---|
| **Oligo #1** | gRNA detection by qPCR Fw | AGCTAAGCTGGAAACAGCA |
| **Oligo #2** | gRNA detection by qPCR Rev | CGACTCGGTGCCACTTT |
| **Oligo #3** | gRNA detection probe | 5'-FAM-AAGGCTAGT/ZEN/CCGTTATCAACTTGAA-3'-IABKFQ |
| **oKK145** | RT primer for gRNA | TTTTTCAAGTTGATAACGGACTAGCC |
| **oKK147** | PCR Rev for gRNA | CCTCTCTATGGGCAGTCGGTGATGACTAGCCTTATTTAAACTTGCTATG |
| **Oligo #4** | RT primer for MS2-gRNA | AAAGCACCGACTCGGTGCCAC |
| **Oligo #5** | PCR Rev for MS2-gRNA | CCTCTCTATGGGCAGTCGGTGATGCCAAGTTGATAACGGACTAGCCTT |
| **LNA-TSO** | template switching oligo | AAGCAGTGGTATCAACGCAGAGTACrGrG+G |
| **oKK120** | qPCR Fw for NGS sample | CCATCTCATCCCTGCGTGTCTCC |
| **oKK121** | qPCR Rev for NGS sample | CCTCTCTATGGGCAGTCGGTGAT |
| **oKK29** | subcloning pKK47 Fw | ATTAGGCCTCTGAGGCCACCATGGCGGTGGAAGGAGGAATGAAATGTGTGAAG |
| **oKK103** | subcloning pKK47 Rev | ATTAGGCCTGACAGGCCTTACGCGTAGATGCCGGAGTTTGCTGCG |
| **oKK109** | subcloning pKK49 Fw | GTCTTCGAGAAGACCTGTTTAAGAGCTAAGCCAAC |
| **oKK105** | subcloning pKK49 Rev | ATTACTCGAGAAAAAAAGCACCGACTCGGTGCCACTTGG |
| **oKK127** | subcloning pKK59 Fw | ATTAGGCCTCTGAGGCCAGCTTGCCACCATGGCGGTG |
| **oKK128** | subcloning pKK59 Rev | ATTAGGCCTGACAGGCCGCTAGCTAATGGTACCGGACCCGCCTCCGCCGAATTC |
| **oKK129** | subcloning pKK60 Fw | ATTAGGTACCGACAAGAAGTACAGCATCGGCCTG |
| **oKK130** | subcloning pKK60 Rev | ATTAGCTAGCCTAGTCGCCTCCCAGCTGAGACAG |
| **oRyo004** | subcloning pRK300 Fw | ATTAGGCCTCTGAGGCCACCATGGACAAGAAGTACAGCATCGG |
| **oRyo005** | subcloning pRK300 Rev | TAATGGCCTGACAGGCCTACTTATCGTCATCGTCTTTGTAATCTTTCTTCTTCTTAGCCTGTCCAG |
| **oKK206** | subcloning pKK89 Fw | TTGGAAAGAGGGTCCCCCTGTAGGTTTAAGAGC |
| **oKK207** | subcloning pKK89 Rev | TTAGCTCTTAAACCTACAGGGGGACCCTCTTTCCAACAAG |

| | | |
|---|---|---|
| **oKK208** | subcloning pKK90 Fw | TTGGAACAAGTTCCAATTTGTATGTTTAAGAGC |
| **oKK209** | subcloning pKK90 Rev | TTAGCTCTTAAACATACAAATTGGAACTTGTTCCAACAAG |
| **oKK531** | subcloning pKK209 Fw | TTGGCACATGAAGCTGTACATGGGTTTAAGAGC |
| **oKK532** | subcloning pKK209 Rev | TTAGCTCTTAAACCCATGTACAGCTTCATGTGCCAACAAG |
| **oKK533** | subcloning pKK210 Fw | TTGGGCGAAGGCAAGCCCTACGAGTTTAAGAGC |
| **oKK534** | subcloning pKK210 Rev | TTAGCTCTTAAACTCGTAGGGCTTGCCTTCGCCCAACAAG |
| **oKK600** | subcloning pKK104 Fw | ATTAGGCCTCTGAGGCCACCATGCCGGTCAAAGGAGGCACCAAG |
| **oKK621** | subcloning pKK104 Rev | ATTAGGCCTGACAGGCCGCTAGCTAATGGTACCGGACCCGCCTCCGCCGAATTCCAAGACCATCTCGCGGTTCCTGCG |
| **oKK236** | subcloning pKK108 Fw | ATTAGGTACCATGGTCTTCACACTCGAAGATTTCG |
| **oKK237** | subcloning pKK108 Rev | ATTAGCTAGCTTACGCCAGAATGCGTTCGC |
| **oRK164** | subcloning pRK397 Fw | ATTAGGTACCGTGAGCAAGGGCGAGGCAGT |
| **oRK165** | subcloning pRK397 Rev | ATTAGCTAGCCTACTTGTACAGCTCGTCCATGCCG |
| **oKK334** | subcloning pKK151 Fw | ATTAGGTACCAGCAAAGGAGAAGAACTTTTCAC |
| **oKK335** | subcloning pKK151 Rev | ATTAGCTAGCCTATTTGTAGAGCTCATCCATGCC |
| **oKK370** | qPCR OSBP Fw | GATCCATCAGGAAAAGTCCAC |
| **oKK371** | qPCR OSBP Rev | CAGTGCCACTTTCCCAAGCA |
| **oKK362** | qPCR FASN Fw | CGCGTGGCCGGCTACTCCTAC |
| **oKK363** | qPCR FASN Rev | CGGCTGCCACACGCTCCTCT |
| **oKK382** | qPCR TMED10 Fw | GAGATGCGTGATACCAACGA |
| **oKK383** | qPCR TMED10 Rev | TTCTTGGCCTTGAAGAAGCG |
| **oKK368** | qPCR GOLGA2 Fw | ACGGATCAGTTGGAAGAAGAAA |
| **oKK369** | qPCR GOLGA2 Rev | GGATCCCTATGGTCTGAATGTG |
| **oKK388** | qPCR PTPN23 Fw | GCCAGCTGTGAAGAAGTTTGT |
| **oKK389** | qPCR PTPN23 Rev | ACAGCCCTCAAAGTC TCGTG |
| **oKK402** | qPCR CAB39 Fw | CACGTTTTTAAGGTGTTTGTAGCC |
| **oKK403** | qPCR CAB39 Rev | ATCCTCCGTCCTGTCGTTCTG |
| **oKK392** | qPCR VPS28 Fw | TTGTTCCCAGGGGCTCCTAT |
| **oKK393** | qPCR VPS28 Rev | ATCTTCCCGACCGCGAG |
| **oKK376** | qPCR PI4KA Fw | GCCTGGAGCATCTCTCCCTA |
| **oKK377** | qPCR PI4KA Rev | AGGCACATCACTAACGGCTC |
| **GAPDH 2F** | qPCR GAPDH Fw | TCCCTGAGCTGAACGGGAAG |
| **GAPDH 2R** | qPCR GAPDH Rev | GGAGGAGTGGGTGTCGCTGT |
| **Oligo #6-#17*** | PCR Fw for gRNA | CCATCTCATCCCTGCGTGTCTCCGACTCAG(**BCD**)AAGCAGTGGTATCAACGCAGAGT |
| **Oligo #18** | PCR Fw for MS2-gRNA library | CCATCTCATCCCTGCGTGTCTCCGACTCAGTACCAAGATCGATGCACAAAAGGAAACTCACCCT |

**Supplementary Table 2.** Oligonucleotides used in this study. The barcode sequences (BCD) of Oligo #6-#17 are listed in Supplementary Table 3. rG, Riboguanosine. +G, Locked guanosine.

Barcode sequences

| Name | BCD (5' -> 3') |
|---|---|
| **Oligo #6** | CTGCAAGTTCGAT |
| **Oligo #7** | TTCGTGATTCGAT |
| **Oligo #8** | TTCCGATAACGAT |
| **Oligo #9** | TGAGCGGAACGAT |
| **Oligo #10** | CTGACCGAACGAT |
| **Oligo #11** | TCCTCGAATCGAT |
| **Oligo #12** | TAGGTGGTTCGAT |
| **Oligo #13** | TCTAACGGACGAT |
| **Oligo #14** | TGCCACGAACGAT |
| **Oligo #15** | AACCTCATTCGAT |
| **Oligo #16** | CCTGAGATACGAT |
| **Oligo #17** | TTACAACCTCGAT |

Correspondence table of NGS samples and oligos used for tagging PCR

| Sample | CD63_CIBER | CD9_CIBER |
|---|---|---|
| **ACOC_cell_rep. 1_Cas9+** | Oligo #14 | Oligo #14 |
| **ACOC_cell_rep. 1_Cas9-** | Oligo #15 | Oligo #15 |
| **ACOC_cell_rep. 2_Cas9+** | Oligo #16 | Oligo #16 |
| **ACOC_cell_rep. 2_Cas9-** | Oligo #17 | Oligo #17 |
| **ACOC_sEVs_rep. 1_Cas9+** | Oligo #14 | Oligo #14 |
| **ACOC_sEVs_rep. 1_Cas9-** | Oligo #15 | Oligo #15 |
| **ACOC_sEVs_rep. 2_Cas9+** | Oligo #16 | Oligo #16 |
| **ACOC_sEVs_rep. 2_Cas9-** | Oligo #17 | Oligo #17 |
| **DTKP_cell_rep. 1_Cas9+** | Oligo #6 | Oligo #14 |
| **DTKP_cell_rep. 1_Cas9-** | Oligo #7 | Oligo #15 |
| **DTKP_cell_rep. 2_Cas9+** | Oligo #8 | Oligo #16 |
| **DTKP_cell_rep. 2_Cas9-** | Oligo #9 | Oligo #17 |
| **DTKP_sEVs_rep. 1_Cas9+** | Oligo #10 | Oligo #14 |
| **DTKP_sEVs_rep. 1_Cas9-** | Oligo #11 | Oligo #15 |
| **DTKP_sEVs_rep. 2_Cas9+** | Oligo #12 | Oligo #16 |
| **DTKP_sEVs_rep. 2_Cas9-** | Oligo #13 | Oligo #17 |
| **PROT_cell_rep. 1_Cas9+** | Oligo #14 | Oligo #14 |
| **PROT_cell_rep. 1_Cas9-** | Oligo #15 | Oligo #15 |
| **PROT_cell_rep. 2_Cas9+** | Oligo #16 | Oligo #16 |
| **PROT_cell_rep. 2_Cas9-** | Oligo #17 | Oligo #17 |
| **PROT_sEVs_rep. 1_Cas9+** | Oligo #14 | Oligo #14 |
| **PROT_sEVs_rep. 1_Cas9-** | Oligo #15 | Oligo #15 |
| **PROT_sEVs_rep. 2_Cas9+** | Oligo #16 | Oligo #16 |
| **PROT_sEVs_rep. 2_Cas9-** | Oligo #17 | Oligo #17 |
| **TMMO_cell_rep. 1_Cas9+** | Oligo #14 | Oligo #14 |
| **TMMO_cell_rep. 1_Cas9-** | Oligo #15 | Oligo #15 |
| **TMMO_cell_rep. 2_Cas9+** | Oligo #16 | Oligo #16 |
| **TMMO_cell_rep. 2_Cas9-** | Oligo #17 | Oligo #17 |
| **TMMO_sEVs_rep. 1_Cas9+** | Oligo #14 | Oligo #14 |
| **TMMO_sEVs_rep. 1_Cas9-** | Oligo #15 | Oligo #15 |

| | | |
|---|---|---|
| **TMMO_sEVs_rep. 2_Cas9+** | Oligo #16 | Oligo #16 |
| **TMMO_sEVs_rep. 2_Cas9-** | Oligo #17 | Oligo #17 |

**Supplementary Table 3.** Barcode information of NGS primers.

| **Name** | **Source** | **Product name** |
|---|---|---|
| **siOSBP#1** | iDT | DsiRNA, 2 nmol (hs.Ri.OSBP.13.1) |
| **siOSBP#2** | iDT | DsiRNA, 2 nmol (hs.Ri.OSBP.13.2) |
| **siOSBP#3** | iDT | DsiRNA, 2 nmol (hs.Ri.OSBP.13.3) |
| **siTMED10#1** | iDT | DsiRNA, 2 nmol (hs.Ri.TMED10.13.1) |
| **siTMED10#2** | iDT | DsiRNA, 2 nmol (hs.Ri.TMED10.13.2) |
| **siTMED10#3** | iDT | DsiRNA, 2 nmol (hs.Ri.TMED10.13.3) |
| **siGOLGA2#1** | iDT | DsiRNA, 2 nmol (hs.Ri.GOLGA2.13.1) |
| **siGOLGA2#2** | iDT | DsiRNA, 2 nmol (hs.Ri.GOLGA2.13.2) |
| **siGOLGA2#3** | iDT | DsiRNA, 2 nmol (hs.Ri.GOLGA2.13.3) |
| **siCtrl for iDT siRNA** | iDT | Negative Control DsiRNA, 1 nmol |
| **siFASN#1** | BIONEER | FASN, GENE_ID 2194, AccuTarget Genome-wide Predesigned siRNA #1 |
| **siFASN#2** | BIONEER | FASN, GENE_ID 2194, AccuTarget Genome-wide Predesigned siRNA #2 |
| **siFASN#3** | BIONEER | FASN, GENE_ID 2194, AccuTarget Genome-wide Predesigned siRNA #3 |
| **siPTPN23#1** | BIONEER | PTPN23, GENE_ID 25930, AccuTarget Genome-wide Predesigned siRNA #1 |
| **siPTPN23#2** | BIONEER | PTPN23, GENE_ID 25930, AccuTarget Genome-wide Predesigned siRNA #2 |
| **siPTPN23#3** | BIONEER | PTPN23, GENE_ID 25930, AccuTarget Genome-wide Predesigned siRNA #3 |
| **siCAB39#1** | BIONEER | CAB39, GENE_ID 51719, AccuTarget Genome-wide Predesigned siRNA #1 |
| **siCAB39#2** | BIONEER | CAB39, GENE_ID 51719, AccuTarget Genome-wide Predesigned siRNA #2 |
| **siCAB39#3** | BIONEER | CAB39, GENE_ID 51719, AccuTarget Genome-wide Predesigned siRNA #3 |
| **siVPS28#1** | BIONEER | VPS28, GENE_ID 51160, AccuTarget Genome-wide Predesigned siRNA #1 |
| **siVPS28#2** | BIONEER | VPS28, GENE_ID 51160, AccuTarget Genome-wide Predesigned siRNA #2 |
| **siVPS28#3** | BIONEER | VPS28, GENE_ID 51160, AccuTarget Genome-wide Predesigned siRNA #3 |
| **siPI4KA#1** | BIONEER | PI4KA, GENE_ID 5297, AccuTarget Genome-wide Predesigned siRNA #1 |
| **siPI4KA#2** | BIONEER | PI4KA, GENE_ID 5297, AccuTarget Genome-wide Predesigned siRNA #2 |
| **siPI4KA#3** | BIONEER | PI4KA, GENE_ID 5297, AccuTarget Genome-wide Predesigned siRNA #3 |
| **siCtrl for BIONEER siRNA** | BIONEER | AccuTarget Negative Control siRNA |

**Supplementary Table 4.** Sources of siRNA used in this study.

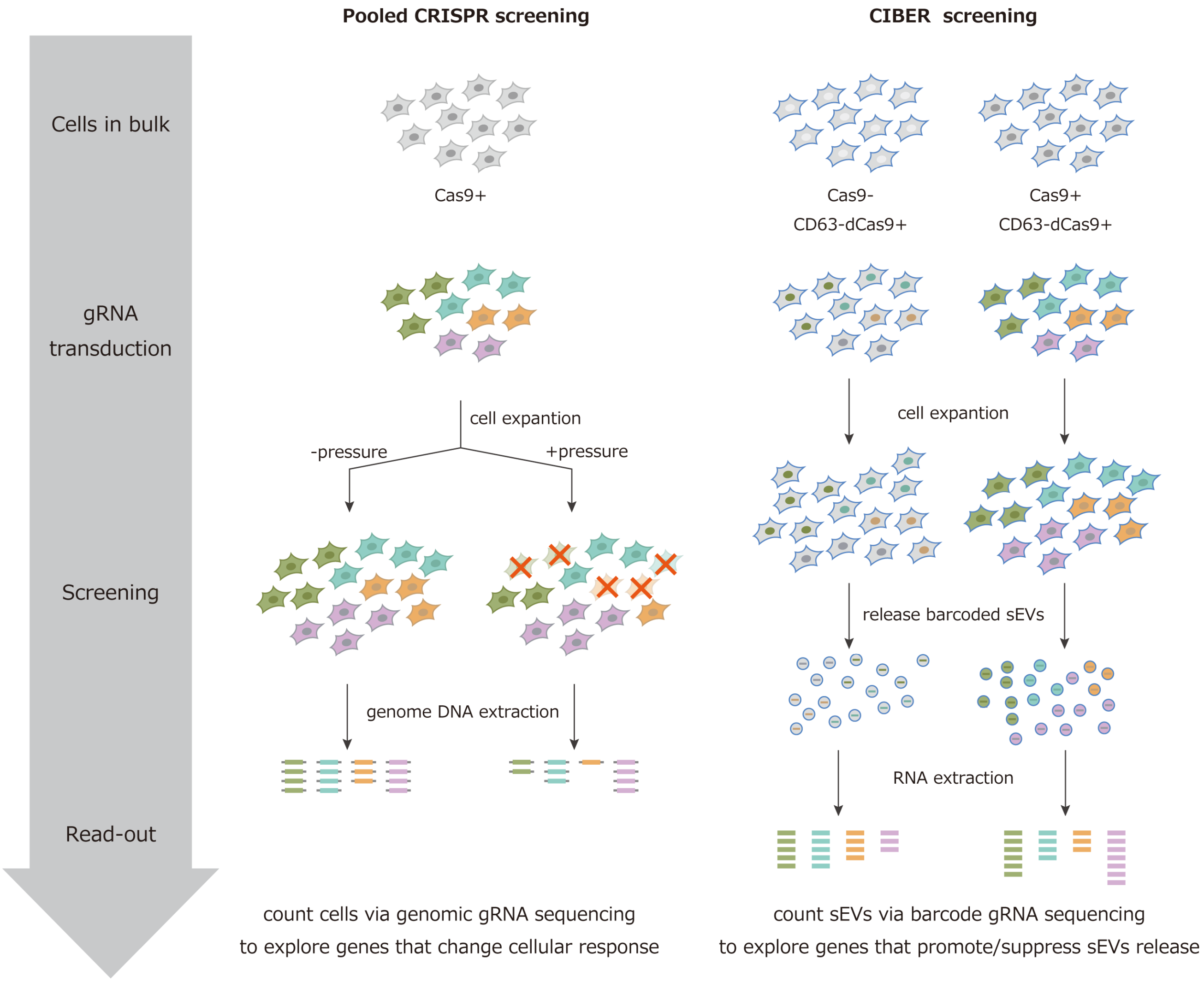


**Supplementary Figure 1** | Comparison of conventional pooled CRISPR screening and CIBER screening. In CIBER screening, transcribed gRNA is used as a barcode while the gRNA expression cassette in the genome is used as a barcode in normal CRISPR screening.

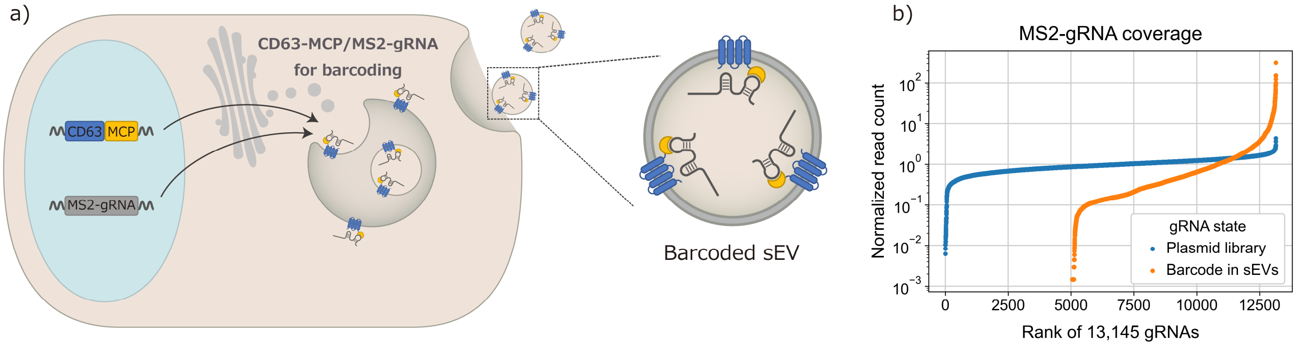


**Supplementary Figure 2 |** Description and performance of MS2-based gRNA loading system. (a) Strategy to barcode sEVs using CD63-MS2 coat protein (MCP) and gRNAs bearing MCP-binding motif (MS2-gRNA). Transcribed MS2-gRNA binds to CD63-fused MCP and is actively loaded into sEVs. (b) Coverage of MS2-gRNA. NGS samples were prepared from a plasmid library consisting of 13,145 MS2-gRNAs (see Supplementary Methods for construction) or RNAs extracted from sEVs released from cells expressing CD63-MCP and transduced with the library. Read counts for every single gRNA were divided by the total count of the sample, then multiplied by the number of gRNAs in the library (13,145) to calculate normalized read count so that the mean value is 1. The barcoding efficacy was not as high as with the CD63-dCas9-based system.

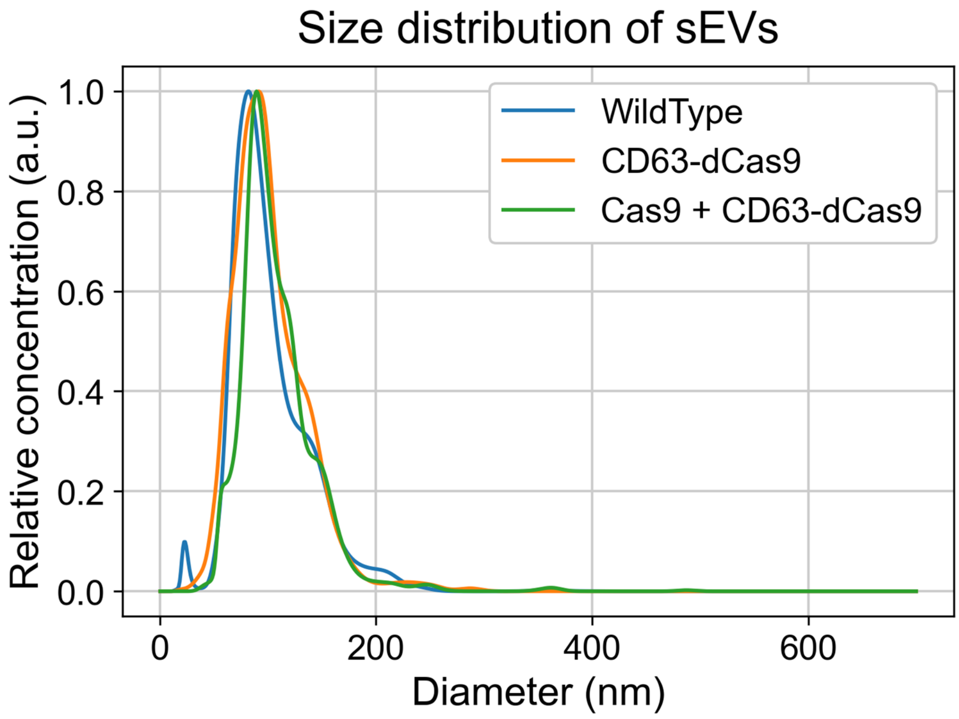


**Supplementary Figure 3 |** Results of nanoparticle tracking analysis (NTA) for sEVs isolated from culture media (CM) of HEK293T cells expressing each component. The concentration and size distribution of sEVs are measured by NanoSight LM10. The lines show averaged concentration from 3 sequential measurements of each sample.

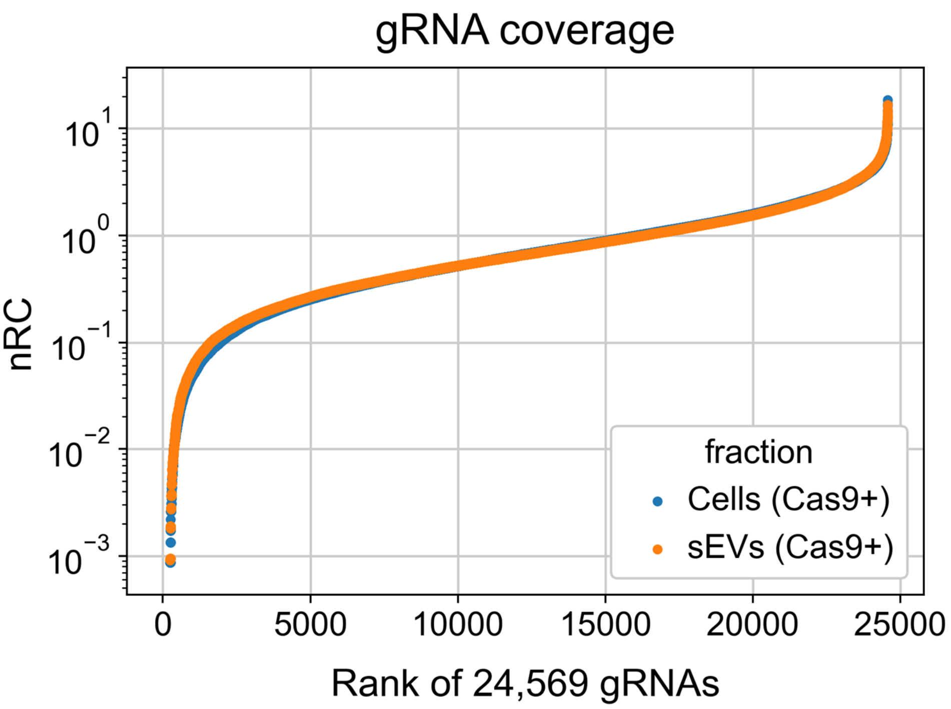


**Supplementary Figure 4 |** Confirmation that co-expression of Cas9 has little effect on coverage of gRNAs in sEVs. Cells expressing Cas9 and CD63-dCas9 were transduced with a library of 24,569 gRNAs. RNAs extracted from cells and sEVs released from them were processed for NGS read-out.

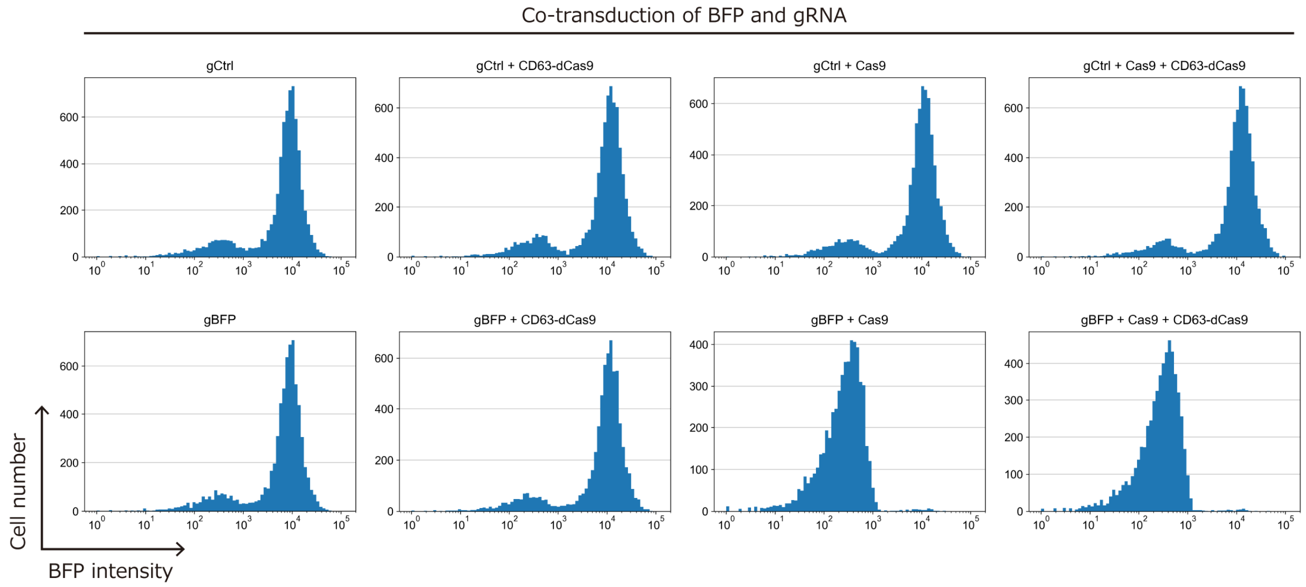


**Supplementary Figure 5 |** Confirmation that CD63-dCas9 expression has little effect on Cas9-induced gene KO. Flow cytometry analyses of HEK293T cells transduced with BFP and either control gRNA (gCtrl, pKK90) or BFP-targeting gRNA (gBFP, pKK209) are shown. BFP-derived fluorescence disappeared in cells expressing Cas9 and gBFP regardless of the expression of CD63-dCas9.

(a)

| cat. # | symbol | Description | Genes targeted | gRNAs |
|---|---|---|---|---|
| 101926 | ACOC | Apoptosis and cancer | 3,015 | 31,324 |
| 101927 | DTKP | Drug targets, kinases, phosphatases | 2,333 | 24,569 |
| 101930 | PROT | Proteostasis | 2,927 | 30,403 |
| 101931 | TMMO | Trafficking, mitochondrial, motility | 2,252 | 23,615 |

(b)

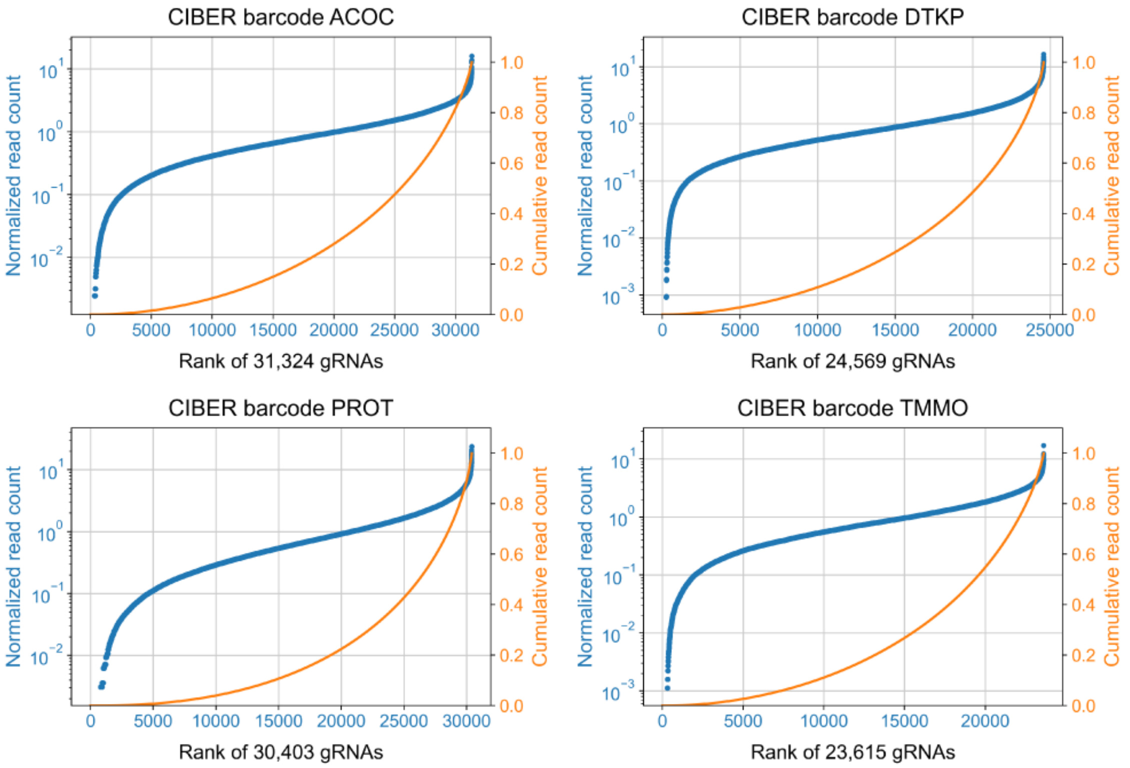


**Supplementary Figure 6 |** gRNA libraries used in this study and their barcoding performance in sEVs. (a) The four listed sub-pool libraries of Bassik deletion libraries[8] covering about half the genome in total were used. The spacer sequences are available in Supplementary data 7. (b) Data showing the high sEV barcoding efficacy using each gRNA library with HEK293T cells expressing CD63-dCas9 and Cas9 (the blue line obtained with the DTKP library is the same as the orange line in Supplementary Fig. 4).

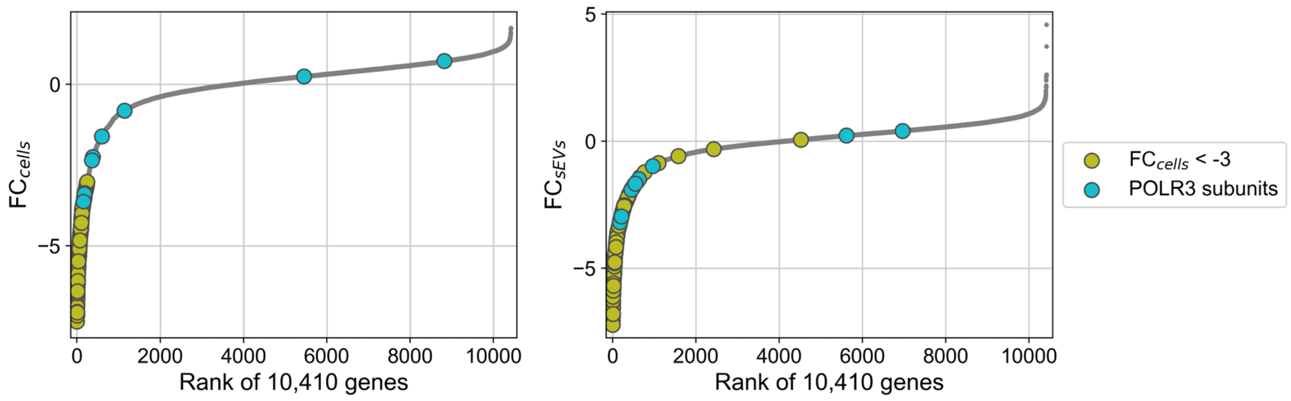


**Supplementary Figure 7 |** Evaluation of the effect of cellular gRNA level on gRNA amount in sEVs. The median value of z-normalized $\log_2$(Fold-change) of barcode gRNAs grouped by target gene in cells ($FC_{cells}$) and sEVs ($FC_{sEVs}$) are shown. gRNAs targeting POLR3 subunits (regulating gRNA transcription) as well as gRNAs targeting genes showing $FC_{cells}$ less than -3 are highlighted. The results indicate that the sEV gRNA level is highly dependent on the cellular gRNA level, showing the necessity of taking this into consideration for extracting true sEV release regulators.

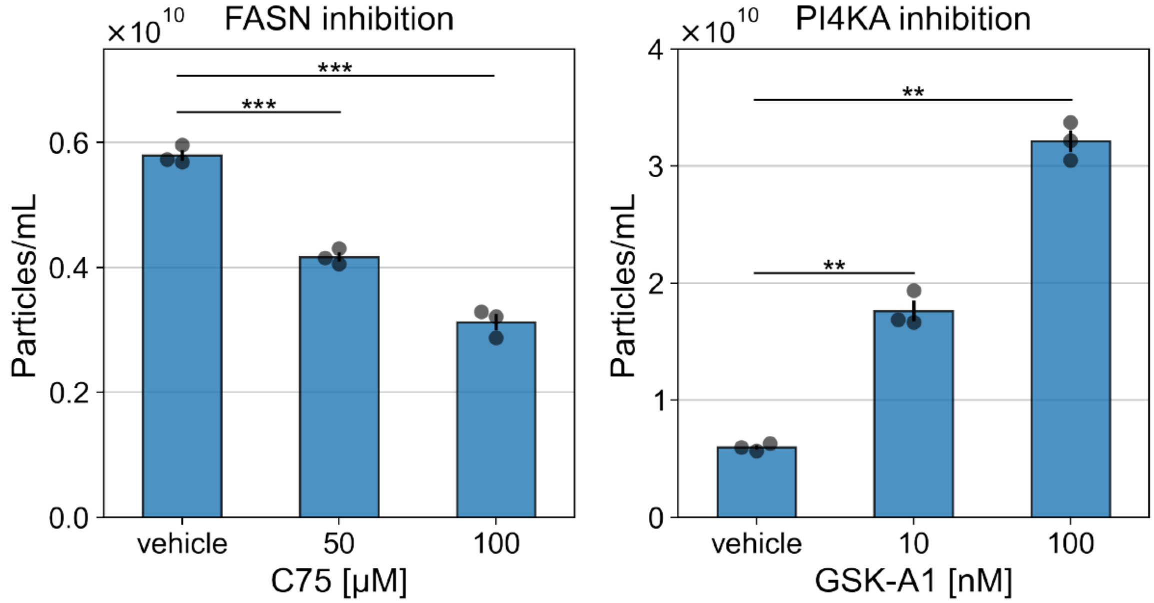


**Supplementary Figure 8 |** Concentration of sEVs released from HEK293T cells treated with C75 (FASN inhibitor) GSK-A1 (PI4KA inhibitor). NTA histograms in Fig. 3d, e were converted to bar graphs showing the particle numbers. Error bars represent ± SEM of biological replicates (n = 3). p: two-tailed Welch's t-test. **$p < 0.005$, ***$p < 0.0005$

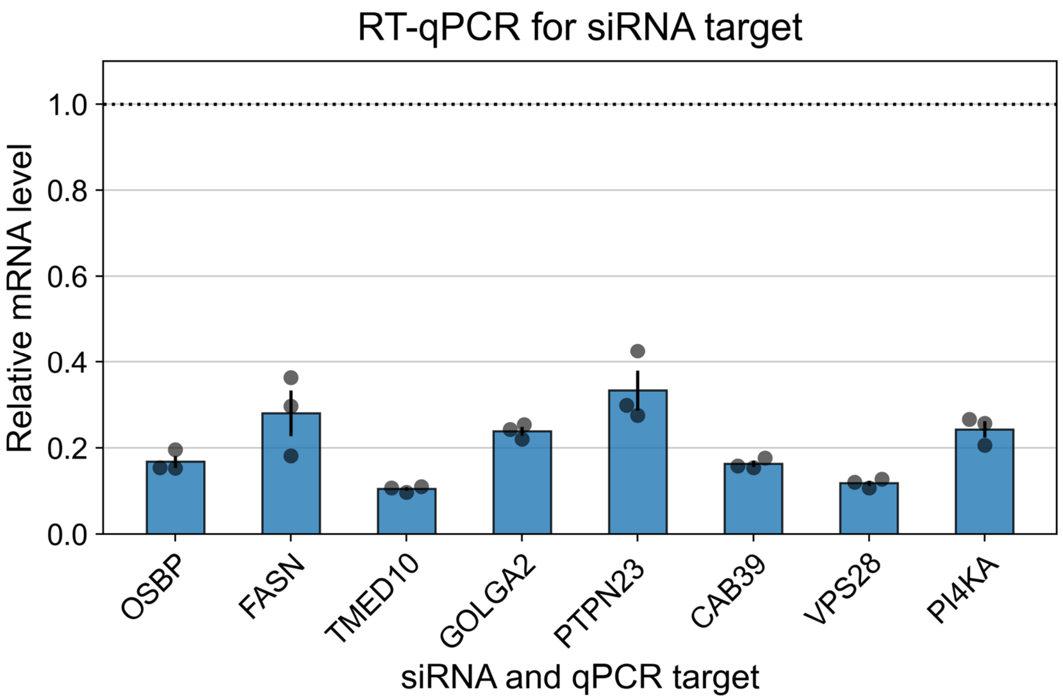


**Supplementary Figure 9 |** Confirmation of gene knockdown by siRNA used for validation in Figure 3f. Cells were transfected with siRNA targeting one of the hit genes and incubated for 24 hours before lysis for qPCR quantification. Relative mRNA level was calculated using the $2^{-\Delta\Delta Ct}$ method with GAPDH as an internal control[12]. Error bars represent ± SEM of biological replicates (n = 3).

| Rank | Gene | Rank | Gene | Rank | Gene | Rank | Gene | Rank | Gene | Rank | Gene | Rank | Gene | Rank | Gene | Rank | Gene | Rank | Gene |
|---|---|---|---|---|---|---|---|---|---|---|---|---|---|---|---|---|---|---|---|
| 1 | CD9 | 101 | ATP6V1A | 201 | CLIC4 | 301 | H2AFX | 401 | PCBP2 | 501 | DLG1 | 601 | SPTBN1 | 701 | CORO1C | 801 | HIST1H2AG | 901 | SERPING1 |
| 2 | ACTB | 102 | PRDX2 | 202 | EEF1G | 302 | RPLP2 | 402 | WARS | 502 | RPS25 | 602 | RPL28 | 702 | CLDN4 | 802 | HIST1H2AI | 902 | TOM1L2 |
| 3 | CD81 | 103 | ACLY | 203 | TXN | 303 | SERPINA1 | 403 | HIST2H2BF | 503 | RPL18 | 603 | LYPLA1 | 703 | HIST1H2AD | 803 | KRT12 | 903 | SYPL1 |
| 4 | HSPA8 | 104 | PROM1 | 204 | CCT3 | 304 | XRCC5 | 404 | ATP6V1E1 | 504 | RPL11 | 604 | HIST1H1D | 704 | LYPLA2 | 804 | VARS | 904 | CORO1B |
| 5 | CD63 | 105 | UBA1 | 205 | PSMA5 | 305 | JUP | 405 | SLC12A1 | 505 | PTPRJ | 605 | RPL35A | 705 | ATP5A1 | 805 | PSMC6 | 905 | ACE2 |
| 6 | GAPDH | 106 | BROX | 206 | RRAS | 306 | HIST1H2BG | 406 | RAB21 | 506 | RPL10A | 606 | GPRC5C | 706 | PPP2R1B | 806 | ALDH9A1 | 906 | TAOK1 |
| 7 | PKM | 107 | YWHAH | 207 | SLC16A1 | 307 | PDIA3 | 407 | PTGR1 | 507 | PCNA | 607 | EPS8L1 | 707 | PSMD5 | 807 | RHOB | 907 | PTP4A1 |
| 8 | ANXA2 | 108 | RAN | 208 | SLC1A5 | 308 | POTEF | 408 | AP2A1 | 508 | ACOT7 | 608 | HIST1H2BN | 708 | TXNRD1 | 808 | AP2M1 | 908 | TWF2 |
| 9 | SDCBP | 109 | TUBA1B | 209 | HSP90B1 | 309 | MYL12B | 409 | IMPDH2 | 509 | KRT6A | 609 | LTF | 709 | RAB22A | 809 | RHEB | 909 | RACGAP1 |
| 10 | HSP90AA1 | 110 | EPCAM | 210 | ARF4 | 310 | GANAB | 410 | SLC1A4 | 510 | MYO1B | 610 | PACSIN3 | 710 | ATP1A2 | 810 | FARSA | 910 | UPF1 |
| 11 | ANXA5 | 111 | ACTN1 | 211 | PDCD10 | 311 | YES1 | 411 | ABHD14B | 511 | FKBP4 | 611 | RAB15 | 711 | HIST1H2BA | 811 | QDPR | 911 | ERP44 |
| 12 | LDHA | 112 | FLOT2 | 212 | RPS16 | 312 | MYO1D | 412 | EIF5A | 512 | PLXNB2 | 612 | PSMD13 | 712 | C11orf54 | 812 | HPGD | 912 | CYBRD1 |
| 13 | HSPA1A | 113 | PSMA7 | 213 | HIST1H2AE | 313 | RHOG | 413 | HINT1 | 513 | DARS | 613 | DPP3 | 713 | PTPRF | 813 | KIF23 | 913 | HNRNPU |
| 14 | YWHAE | 114 | RAB14 | 214 | AKR1A1 | 314 | ALDOC | 414 | HRAS | 514 | NAMPT | 614 | KIF5B | 714 | EIF2S1 | 814 | GSS | 914 | EIF5 |
| 15 | YWHAZ | 115 | RAB5A | 215 | ACTBL2 | 315 | AARS | 415 | RPS24 | 515 | CA2 | 615 | RPL27 | 715 | ATP1A3 | 815 | GSTM2 | 915 | CAV1 |
| 16 | PDCD6IP | 116 | CAND1 | 216 | ARPC2 | 316 | RAB13 | 416 | DDT | 516 | EIF4A2 | 616 | RPL23 | 716 | PABPC1 | 816 | EPHB3 | 916 | RAB25 |
| 17 | ENO1 | 117 | CCT5 | 217 | ITGA3 | 317 | ACTA2 | 417 | GPRC5A | 517 | GFPT1 | 617 | TMPRSS2 | 717 | NAPG | 817 | HIST1H2AA | 917 | UBE2K |
| 18 | ALB | 118 | TAGLN2 | 218 | LSR | 318 | NT5E | 418 | RPL4 | 518 | VPS35 | 618 | UMOD | 718 | RAB18 | 818 | KRT3 | 918 | COL18A1 |
| 19 | MSN | 119 | GNAI3 | 219 | RPS8 | 319 | HIST1H2BL | 419 | HIST1H2BF | 519 | KRT6C | 619 | NAP1L1 | 719 | CRK | 819 | 7-Sep | 919 | HCK |
| 20 | EEF1A1 | 120 | RPS27A | 220 | PGD | 320 | CYFIP1 | 420 | SCARB2 | 520 | HPRT1 | 620 | NACA | 720 | EFNB1 | 820 | DOPEY2 | 920 | USP9X |
| 21 | CFL1 | 121 | PSMA4 | 221 | KRT2 | 321 | SOD1 | 421 | PLS3 | 521 | VAMP8 | 621 | PEPD | 721 | FABP5 | 821 | SLK | 921 | TGM2 |
| 22 | TPI1 | 122 | CBR1 | 222 | PSMB2 | 322 | CAPZB | 422 | HEBP1 | 522 | NPEPPS | 622 | APOA1 | 722 | ATP4A | 822 | IGF2R | 922 | FAS |
| 23 | LDHB | 123 | CCT6A | 223 | TF | 323 | RRAS2 | 423 | GGT1 | 523 | EPRS | 623 | HIST1H2BJ | 723 | XPO1 | 823 | CD55 | 923 | COPS8 |
| 24 | ACTN4 | 124 | MUC1 | 224 | HSPA6 | 324 | F11R | 424 | HIST2H2AC | 524 | DAK | 624 | VAMP3 | 724 | GLG1 | 824 | CEACAM5 | 924 | RPS15 |
| 25 | PFN1 | 125 | ACTA1 | 225 | PPP1CA | 325 | S100A11 | 425 | HIST1H2BE | 525 | EIF3M | 625 | ACO1 | 725 | CDK1 | 825 | ENPEP | 925 | NANS |
| 26 | FASN | 126 | MDH1 | 226 | HSPG2 | 326 | HIST1H2BB | 426 | PDXK | 526 | KIF12 | 626 | SARS | 726 | NCKAP1 | 826 | UBC | 926 | CHP1 |
| 27 | ANXA6 | 127 | RAP2B | 227 | SNAP23 | 327 | CHMP1B | 427 | CTNNB1 | 527 | NCL | 627 | SELENBP1 | 727 | KIAA0368 | 827 | CS | 927 | RTN4 |
| 28 | YWHAQ | 128 | A2M | 228 | SRC | 328 | RPS2 | 428 | XRCC6 | 528 | PSMD2 | 628 | HSPH1 | 728 | MYH11 | 828 | PCBD1 | 928 | KIF3A |
| 29 | GNAS | 129 | KRT1 | 229 | RPS3 | 329 | C3 | 429 | S100A8 | 529 | MARCKSL1 | 629 | NUTF2 | 729 | RPS19 | 829 | GLIPR2 | 929 | QPRT |
| 30 | PGK1 | 130 | HLA-A | 230 | CSTB | 330 | CIB1 | 430 | S100A9 | 530 | ACY1 | 630 | PSMB8 | 730 | RAB3A | 830 | GPC1 | 930 | ADSS |
| 31 | HSPA1B | 131 | PRDX5 | 231 | PRSS8 | 331 | B2M | 431 | VPS25 | 531 | EHD3 | 631 | GNAO1 | 731 | BHMT | 831 | CANX | 931 | DCTN2 |
| 32 | ALDOA | 132 | FLNA | 232 | GNAQ | 332 | MVP | 432 | MYH14 | 532 | CAPN2 | 632 | EPHB1 | 732 | CAPN7 | 832 | SLC25A5 | 932 | CSNK2B |
| 33 | HIST1H4I | 133 | ACTG1 | 233 | VCL | 333 | ITGA2 | 433 | P4HB | 533 | STXBP3 | 633 | PAFAH1B2 | 733 | ASAH1 | 833 | MGAM | 933 | ECH1 |
| 34 | FLOT1 | 134 | TUBB4B | 234 | CAPZA2 | 334 | RUVBL2 | 434 | KRT15 | 534 | PA2G4 | 634 | CYFIP2 | 734 | KRT4 | 834 | UPK2 | 934 | SSRP1 |
| 35 | EZR | 135 | SLC9A3R1 | 235 | PSMB1 | 335 | TUBB4A | 435 | H2AFV | 535 | UBE2V1 | 635 | PSMC4 | 735 | LGALS1 | 835 | VAMP7 | 935 | LMNA |
| 36 | LGALS3BP | 136 | PSMA3 | 236 | DPEP1 | 336 | ICAM1 | 436 | KRT5 | 536 | PFKP | 636 | CHMP6 | 736 | C4B | 836 | CA4 | 936 | HUWE1 |
| 37 | HSP90AB1 | 137 | CPNE3 | 237 | ACE | 337 | RPS13 | 437 | STX4 | 537 | VIM | 637 | LAMA5 | 737 | GNG5 | 837 | MAN1A1 | 937 | TOP1 |
| 38 | PPIA | 138 | ANXA7 | 238 | GNAI1 | 338 | CAPNS1 | 438 | KRT17 | 538 | GART | 638 | TTN | 738 | RPL34 | 838 | SERPINA3 | 938 | ARPC5 |
| 39 | ANXA1 | 139 | SLC12A2 | 239 | KRT19 | 339 | VAT1 | 439 | CLTCL1 | 539 | HIST1H2BM | 639 | AK1 | 739 | FRK | 839 | ACP2 | 939 | EIF3F |
| 40 | TUBB | 140 | PSMA6 | 240 | KRT13 | 340 | ASS1 | 440 | PLEC | 540 | CAD | 640 | PAFAH1B1 | 740 | GIPC1 | 840 | ACTG2 | 940 | RPL22 |
| 41 | RAB5C | 141 | VPS37B | 241 | UBB | 341 | KRT16 | 441 | HIST1H2BC | 541 | HYOU1 | 641 | SYNCRIP | 741 | ACTR1B | 841 | VPS26A | 941 | GNPNAT1 |
| 42 | TSG101 | 142 | TUBA1A | 242 | RHOC | 342 | CTNND1 | 442 | DNAJA2 | 542 | PYGL | 642 | LYZ | 742 | FBP1 | 842 | MPST | 942 | SGTA |
| 43 | PRDX1 | 143 | CALM2 | 243 | CALM1 | 343 | ENO3 | 443 | IPO5 | 543 | RNPEP | 643 | LYN | 743 | PHB | 843 | TNPO1 | 943 | CSNK2A1 |
| 44 | CLTC | 144 | CNDP2 | 244 | ANPEP | 344 | MYH10 | 444 | FSCN1 | 544 | PSMC5 | 644 | HIST1H2AH | 744 | EIF4A3 | 844 | KARS | 944 | ANKFY1 |
| 45 | RAP1B | 145 | MFGE8 | 245 | PGLS | 345 | SLC7A5 | 445 | DDAH1 | 545 | USP7 | 645 | CUL4A | 745 | H2AFY | 845 | PRKCSH | 945 | RPL37A |
| 46 | GNB2 | 146 | TUBA4A | 246 | CCT8 | 346 | KRAS | 446 | PLS1 | 546 | PRKAR2A | 646 | HNRNPH1 | 746 | PGM1 | 846 | AP1G1 | 946 | ERP29 |
| 47 | CLIC1 | 147 | UBE2N | 247 | CALM3 | 347 | DNAJA1 | 447 | RPS15A | 547 | UBR4 | 647 | KNG1 | 747 | TUBA3C | 847 | ANP32A | 947 | FH |
| 48 | GPI | 148 | TKT | 248 | PIGR | 348 | APEH | 448 | RAC3 | 548 | SET | 648 | GPRC5B | 748 | MST4 | 848 | CHMP1A | 948 | DDX6 |
| 49 | EEF2 | 149 | NRAS | 249 | ENO2 | 349 | NME2 | 449 | EPHA2 | 549 | QSOX1 | 649 | SLC16A3 | 749 | HIST1H2AK | 849 | PSMD1 | 949 | PSMB7 |
| 50 | YWHAG | 150 | ACTR3 | 250 | MARCKS | 350 | MDH2 | 450 | PSAT1 | 550 | NQO1 | 650 | AHNAK | 750 | MPP5 | 850 | PCMT1 | 950 | UBE2M |
| 51 | RAB7A | 151 | LAMP1 | 251 | MYL6 | 351 | GRB2 | 451 | RPL30 | 551 | RPS10 | 651 | DERA | 751 | ITIH4 | 851 | PSMC2 | 951 | CSNK1A1 |
| 52 | RDX | 152 | KPNB1 | 252 | RPS5 | 352 | CAB39 | 452 | TNIK | 552 | TPP2 | 652 | AP1B1 | 752 | PTK7 | 852 | FKBP1A | 952 | HGS |
| 53 | GNB1 | 153 | ARHGDIA | 253 | APOE | 353 | CSE1L | 453 | CD46 | 553 | RPL24 | 653 | FDPS | 753 | PLOD3 | 853 | RAD23B | 953 | ATP6V0D1 |
| 54 | RAB10 | 154 | TUBB3 | 254 | APRT | 354 | CAPZA1 | 454 | TUBB1 | 554 | PYGB | 654 | TSPAN8 | 754 | HPX | 854 | SEC24C | 954 | RBP5 |
| 55 | YWHAB | 155 | PARK7 | 255 | KRT8 | 355 | PNP | 455 | AP2A2 | 555 | GNB4 | 655 | ARL3 | 755 | GNPDA1 | 855 | GNA14 | 955 | LRP2 |
| 56 | GNAI2 | 156 | CHMP5 | 256 | ANXA3 | 356 | HNRNPK | 456 | TPM3 | 556 | HSPA9 | 656 | SDC4 | 756 | PTP4A2 | 856 | ARHGDIB | 956 | H3F3C |
| 57 | CDC42 | 157 | PSMB3 | 257 | KRT14 | 357 | HIST2H2BE | 457 | BAIAP2L1 | 557 | SDCBP2 | 657 | EGF | 757 | PSME2 | 857 | BLVRA | 957 | TBCA |
| 58 | VCP | 158 | KRT10 | 258 | RNH1 | 358 | GDI1 | 458 | RPS11 | 558 | HIST1H2AC | 658 | PSMD3 | 758 | H2AFZ | 858 | PSMC1 | 958 | PDZK1 |
| 59 | RALA | 159 | CAP1 | 259 | DYNC1H1 | 359 | NAPA | 459 | STIP1 | 559 | PACSIN2 | 659 | GMDS | 759 | GGH | 859 | DNAJC13 | 959 | DDB1 |
| 60 | ATP1A1 | 160 | ARPC4 | 260 | UEVLD | 360 | STK24 | 460 | CD2AP | 560 | PPP2CA | 660 | RAP2A | 760 | MYO6 | 860 | ATP6V0A1 | 960 | RPLP1 |
| 61 | MYH9 | 161 | TCP1 | 261 | RPL12 | 361 | HNRNPA2B1 | 461 | AP2B1 | 561 | KRT76 | 661 | HBB | 761 | CUL4B | 861 | LIN7C | 961 | ILK |
| 62 | CLU | 162 | RAB2A | 262 | RPS18 | 362 | PPA1 | 462 | RAB43 | 562 | NME1-NME2 | 662 | TPP1 | 762 | RHOF | 862 | AHCYL1 | 962 | SLC25A3 |
| 63 | PDCD6 | 163 | EIF4A1 | 263 | ATP5B | 363 | AKR1B1 | 463 | PSMD7 | 563 | KRT24 | 663 | SERPINB6 | 763 | CHMP3 | 863 | C9 | 963 | NOTCH1 |
| 64 | HSPA5 | 164 | TOLLIP | 264 | ST13 | 364 | LAMC1 | 464 | ESD | 564 | PRDX4 | 664 | DDX3X | 764 | SNRPD3 | 864 | PTPRA | 964 | DNPEP |
| 65 | AHCY | 165 | ACTR2 | 265 | RAB6A | 365 | PFKL | 465 | EGFR | 565 | SCRIB | 665 | AK2 | 765 | PRKACA | 865 | XPO7 | 965 | FABP1 |
| 66 | CHMP2A | 166 | PHGDH | 266 | PCBP1 | 366 | PSMB4 | 466 | VTN | 566 | CDH1 | 666 | DDAH2 | 766 | ITGB5 | 866 | TUBA3E | 966 | TUFM |
| 67 | EHD4 | 167 | PPIB | 267 | IDH1 | 367 | RPS7 | 467 | RPL7 | 567 | MARS | 667 | RPL23A | 767 | PTGES3 | 867 | PFAS | 967 | LIN7A |
| 68 | ARF1 | 168 | CCT7 | 268 | EPS8L2 | 368 | RAB3D | 468 | RAB3B | 568 | DCD | 668 | COTL1 | 768 | PGAM2 | 868 | RPL14 | 968 | FTH1 |
| 69 | RAB1A | 169 | SLC2A1 | 269 | DSP | 369 | ARPC3 | 469 | ALDH1A1 | 569 | SNRNP200 | 669 | PARP1 | 769 | GNAL | 869 | MYO1E | 969 | APCS |
| 70 | MYO1C | 170 | PGAM1 | 270 | OLA1 | 370 | TPM4 | 470 | CLDN3 | 570 | VDAC1 | 670 | AKR7A2 | 770 | CP | 870 | YBX1 | 970 | SLC12A7 |
| 71 | IST1 | 171 | ARF6 | 271 | ITGAV | 371 | ALDOB | 471 | THY1 | 571 | PPP1CC | 671 | PSMC3 | 771 | DNM2 | 871 | ALDH7A1 | 971 | COPG1 |
| 72 | RAP1A | 172 | ARF5 | 272 | FLNB | 372 | CRYZ | 472 | ITGB4 | 572 | SLC29A1 | 672 | GCN1L1 | 772 | RPL31 | 872 | EIF3K | 972 | PSMA8 |
| 73 | HSPA2 | 173 | FN1 | 273 | TUBB2A | 373 | SORD | 473 | G6PD | 573 | ACPP | 673 | DCXR | 773 | HNRNPF | 873 | ABCC1 | 973 | CKAP5 |
| 74 | ANXA4 | 174 | HIST1H2AB | 274 | ATIC | 374 | PSME1 | 474 | GSTA1 | 574 | SPR | 674 | RAB27A | 774 | CPM | 874 | ITGB3 | 974 | UPK1A |
| 75 | MIF | 175 | HSPD1 | 275 | UGP2 | 375 | UGDH | 475 | HLA-B | 575 | RARS | 675 | FLNC | 775 | LCK | 875 | TUBA3D | 975 | SKP1 |
| 76 | RAB35 | 176 | PAICS | 276 | GSN | 376 | MME | 476 | ABCB1 | 576 | HIST1H2BK | 676 | DDX39B | 776 | EFHD2 | 876 | SYNGR2 | 976 | EPHB4 |
| 77 | GDI2 | 177 | ADAM10 | 277 | TUBB6 | 377 | TARS | 477 | RPL13 | 577 | PPP1CB | 677 | TUBB8 | 777 | CST3 | 877 | IPO7 | 977 | RPL35 |
| 78 | GSTP1 | 178 | GOT1 | 278 | ACTR1A | 378 | PDIA6 | 478 | KRT18 | 578 | STX3 | 678 | ARPC1B | 778 | HIST2H2AB | 878 | TUBB2B | 978 | UBL3 |
| 79 | DPP4 | 179 | CD44 | 279 | KRT6B | 379 | TGFBI | 479 | APP | 579 | DOCK9 | 679 | EIF3A | 779 | PLSCR1 | 879 | KRT27 | 979 | FTL |
| 80 | RHOA | 180 | WDR1 | 280 | GSTO1 | 380 | THBS1 | 480 | RUVBL1 | 580 | DHX9 | 680 | AHSG | 780 | FGG | 880 | ITCH | 980 | HSPE1 |
| 81 | VPS28 | 181 | MITD1 | 281 | CFL2 | 381 | HNRNPC | 481 | CKB | 581 | CMPK1 | 681 | IARS | 781 | SERPINA5 | 881 | PKP3 | 981 | ITM2B |
| 82 | RAB8B | 182 | BSG | 282 | RPS14 | 382 | CTSD | 482 | RPS20 | 582 | GRHPR | 682 | RSU1 | 782 | H3F3B | 882 | RPL13A | 982 | ERBB2 |
| 83 | TFRC | 183 | KRT9 | 283 | ITGA6 | 383 | TLN1 | 483 | PSMD6 | 583 | EEF1E1 | 683 | SCARB1 | 783 | FGB | 883 | ACSL4 | 983 | ATP2B2 |
| 84 | HSPB1 | 184 | PSMB6 | 284 | CCT4 | 384 | VASP | 484 | LCP1 | 584 | SLC12A3 | 684 | NAPSA | 784 | RPS28 | 884 | NEDD4L | 984 | CALB1 |
| 85 | RAB5B | 185 | RALB | 285 | RPS4X | 385 | DSTN | 485 | PSAP | 585 | IQGAP2 | 685 | RPL19 | 785 | PRKDC | 885 | USP14 | 985 | MYL6B |
| 86 | RAB1B | 186 | RAC2 | 286 | RPL3 | 386 | NCSTN | 486 | RPL15 | 586 | MASP2 | 686 | SNRPD2 | 786 | PSMD12 | 886 | PDIA4 | 986 | PPL |
| 87 | PRDX6 | 187 | TUBA1C | 287 | SLC44A1 | 387 | ATP6V1B1 | 487 | EPHB2 | 587 | LTA4H | 687 | NARS | 787 | LMAN2 | 887 | CARS | 987 | PPP1R7 |
| 88 | ACTC1 | 188 | RAB11B | 288 | KRT7 | 388 | CTNNA1 | 488 | MINK1 | 588 | CPNE1 | 688 | RP2 | 788 | EIF2S3 | 888 | PLCB3 | 988 | MTAP |
| 89 | ANXA11 | 189 | HLA-DRA | 289 | HSPA4 | 389 | RPL7A | 489 | RAB27B | 589 | STX7 | 689 | LASP1 | 789 | DBI | 889 | RPS12 | 989 | SERPINF1 |
| 90 | SFN | 190 | GNA11 | 290 | LAP3 | 390 | MTHFD1 | 490 | HNRNPD | 590 | SND1 | 690 | CAPN5 | 790 | OLFM4 | 890 | ATP6V1H | 990 | S100P |
| 91 | EHD1 | 191 | LAMP2 | 291 | PSMB5 | 391 | AQP2 | 491 | RPL5 | 591 | EIF3B | 691 | H3F3A | 791 | TPM1 | 891 | GBF1 | 991 | GAA |
| 92 | CCT2 | 192 | EEF1A2 | 292 | VPS4B | 392 | XPNPEP2 | 492 | HIST1H2BH | 592 | RPS6 | 692 | OTUB1 | 792 | ABCE1 | 892 | SERINC1 | 992 | PKLR |
| 93 | PEBP1 | 193 | PTGFRN | 293 | NPM1 | 393 | ACAT2 | 493 | MAP4K4 | 593 | PTBP1 | 693 | PICALM | 793 | ETF1 | 893 | TUBA8 | 993 | MPO |
| 94 | HSPA1L | 194 | PSMA1 | 294 | RAB11A | 394 | ATP6V1B2 | 494 | GARS | 594 | SPTAN1 | 694 | ARPC5L | 794 | BANF1 | 894 | CDC37 | 994 | ADSL |
| 95 | RAC1 | 195 | SRI | 295 | EPS8 | 395 | GSTM3 | 495 | CALR | 595 | PPP2R1A | 695 | DLD | 795 | ROCK2 | 895 | ALCAM | 995 | MAT2A |
| 96 | ITGB1 | 196 | STOM | 296 | CAPN1 | 396 | TALDO1 | 496 | 2-Sep | 596 | ACP1 | 696 | STXBP2 | 796 | LARS | 896 | LAMB1 | 996 | NUMA1 |
| 97 | IQGAP1 | 197 | RPSA | 297 | S100A6 | 397 | RPS9 | 497 | UBA52 | 597 | RPL8 | 697 | RPL17 | 797 | CPNE8 | 897 | C4A | 997 | MPP6 |
| 98 | CHMP4B | 198 | RPLP0 | 298 | GNB2L1 | 398 | GSR | 498 | GOLGA7 | 598 | GLO1 | 698 | APOD | 798 | FGA | 898 | CACYBP | 998 | TRIP10 |
| 99 | RAB8A | 199 | LGALS3 | 299 | CHMP2B | 399 | RPS3A | 499 | VIL1 | 599 | PSMD11 | 699 | TJP2 | 799 | TINAGL1 | 899 | RPN2 | 999 | MARK2 |
| 100 | GNA13 | 200 | CNP | 300 | EPB41L2 | 400 | DPYSL2 | 500 | HRSP12 | 600 | SDC1 | 700 | FYN | 800 | GSTA2 | 900 | ADH5 | 1000 | SNX18 |

| Rank |
|---|
| 1-250 |
| 251-500 |
| 501-750 |
| 751-1000 |

| Gene group |
|---|
| Lower hits |
| N.S. |
| Upper hits |

**Supplementary Figure 10 |** Matching of the hit genes of CD63-CIBER and top 1,000 sEVs-associated proteins compiled from the VesiclePedia database[13,14] (related to Fig. 3g). The lower/upper hits in CD63-CIBER screen are colored blue/orange, respectively.

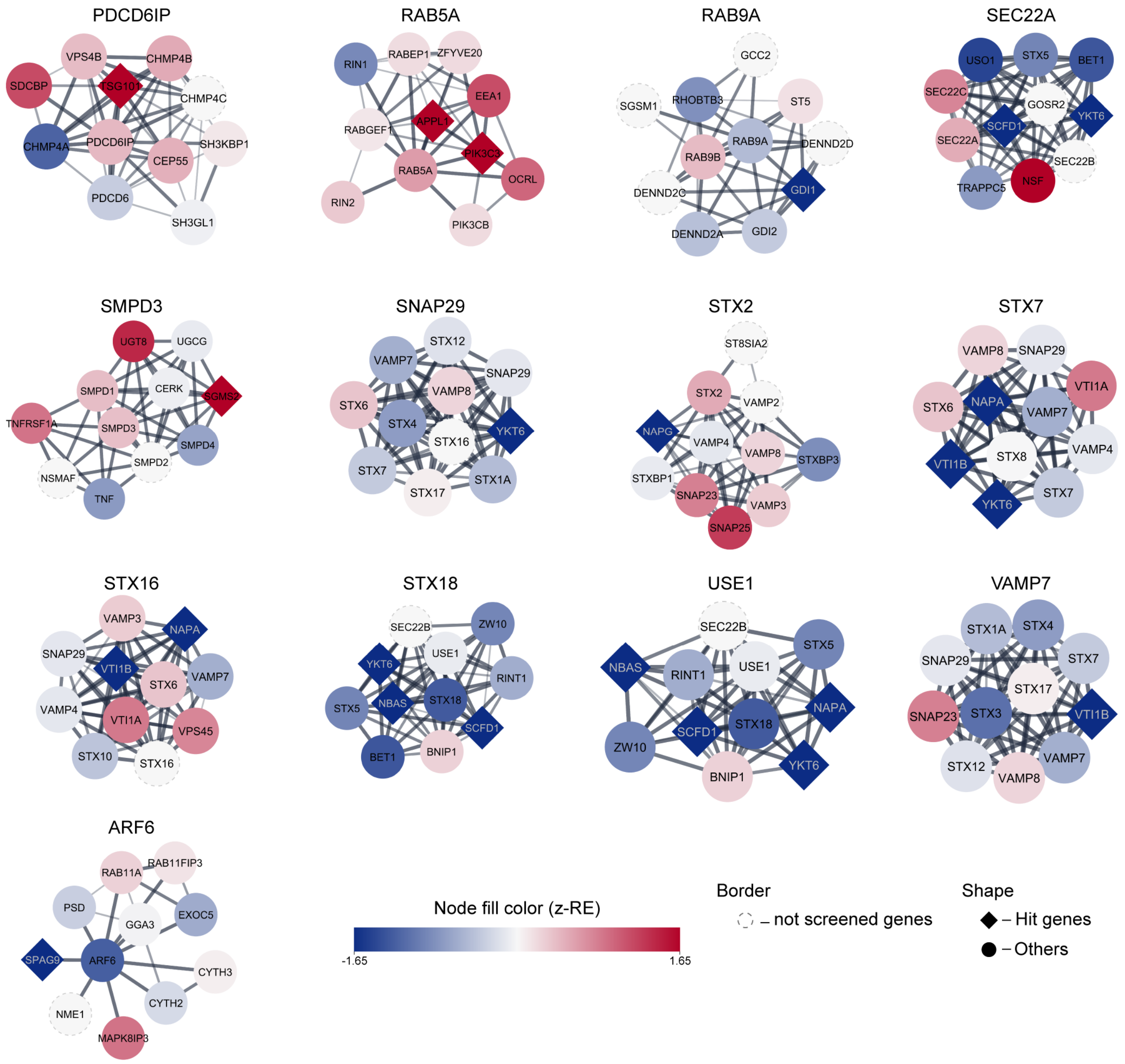


**Supplementary Figure 11 |** Connection of the hit genes in CD63-CIBER with known sEV release regulators (PDCD6IP, RAB5A, RAB9A, SEC22A, SMPD3, SNAP29, STX2, STX7, STX16, STX18, USE1, VAMP7, and ARF6) analyzed by STRING. For each known regulator, the interaction network with functionally connected proteins is displayed under the gene names. Each node represents a gene product, and the node color corresponds to the z-RE. Hit genes are indicated as diamond shapes.

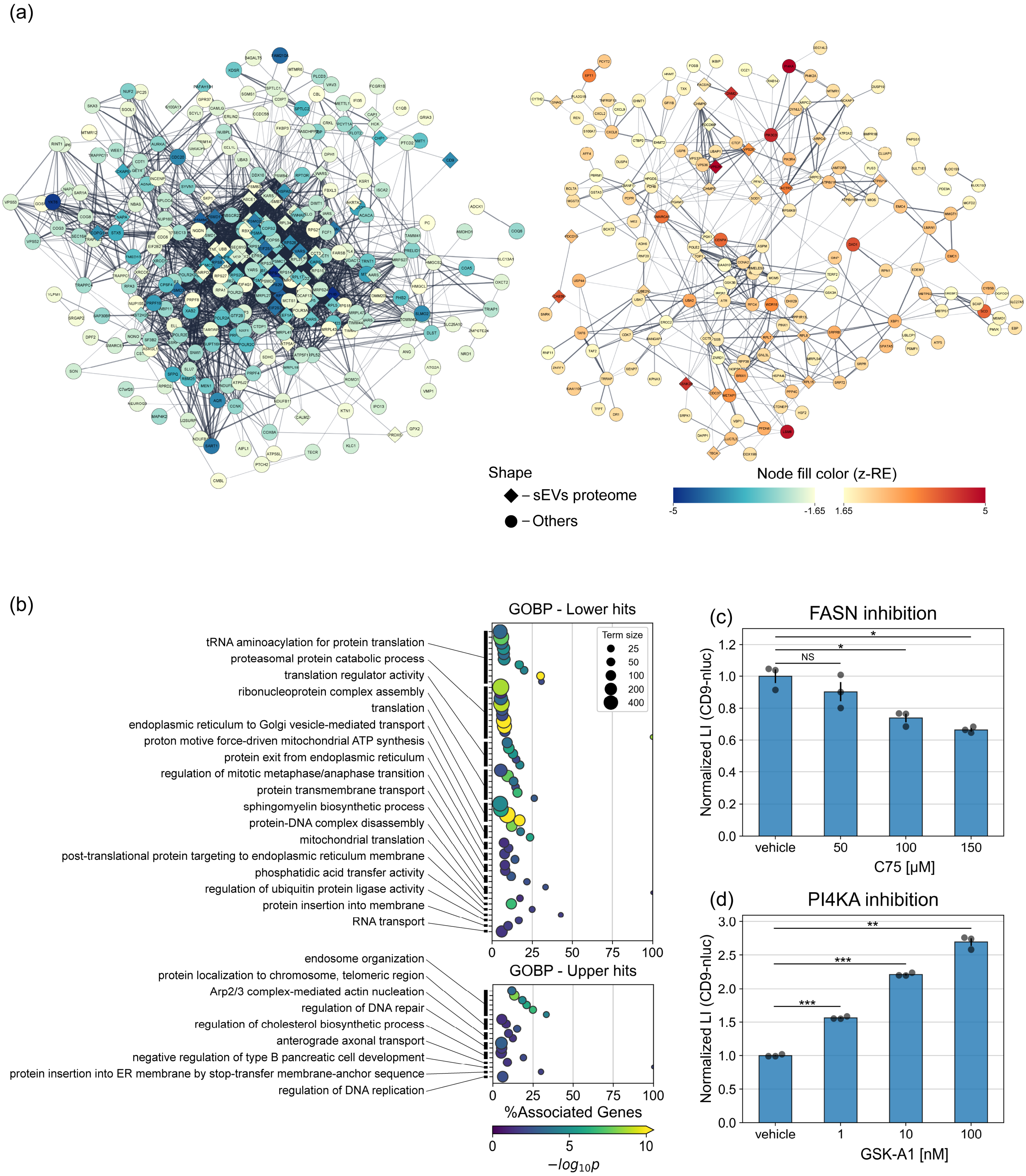


**Supplementary Figure 12 |** Results of CD9-CIBER screening. (a) STRING analysis with lower hits and upper hits of CD9-CIBER screening. Each node represents a gene product, and the node color corresponds to the z-RE. Only nodes connected to at least one other node are shown. The nodes of proteins found in sEVs proteome (Supplementary Fig. 10) are shown as diamond shapes while others are circles. (b) The results of GOBP enrichment analysis. The x-axis represents the gene ratio, which refers to the ratio of lower or upper hits genes to all gene numbers annotated to the term. The circle size indicates the total number of genes annotated to the term. The circle color indicates the -$\log_{10}$(p-value) by Fisher's exact test adjusted by Holm correction. Only terms with adjusted p-value lower than 0.05 are displayed. Terms are grouped based on their kappa score level (0.4) calculated using ClueGO app and represented by the most significant term in each group. (c, d) Validation of the effect of PI4KA and FASN with the CD9-nanoluc (nluc) reporter assay.

HEK293T cells stably expressing CD9-nluc were treated as described in Fig 3b, c. Error bars represent ± SEM of biological replicates (n=3). p: two-tailed Welch's t-test. *p < 0.05, **p < 0.005, ***p < 0.0005, NS, not significant.

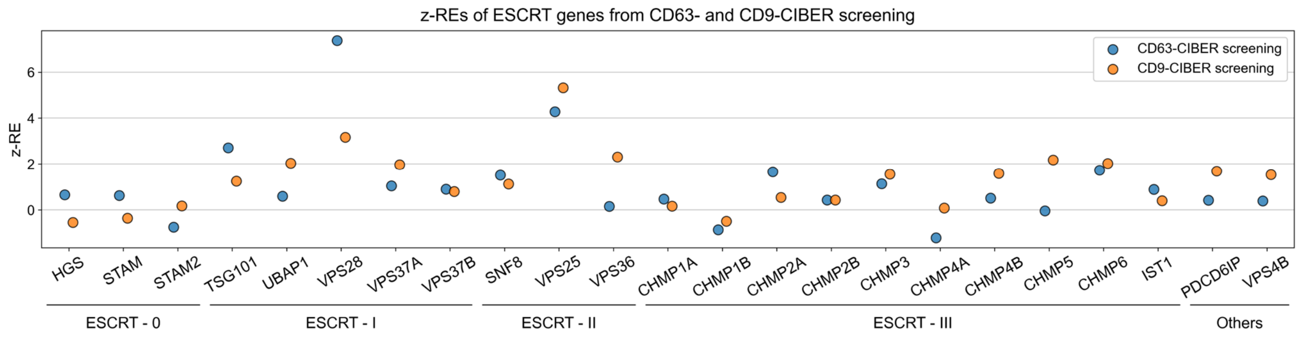


**Supplementary Figure 13 |** z-RE of ESCRT genes obtained from CD63- and CD9-CIBER screening. R = 0.63.

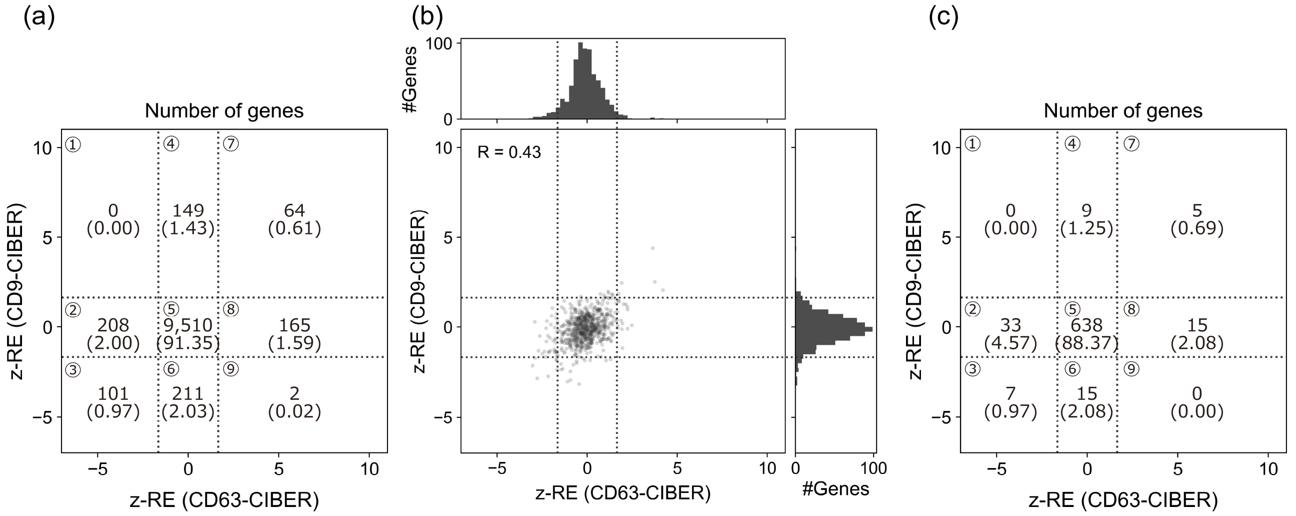


(d)

| Term | Term size | CD63-CIBER | | CD9-CIBER | |
|---|---|---|---|---|---|
| | | #Associated genes | adjp | #Associated genes | adjp |
| NADH dehydrogenase (ubiquinone) activity | 42 | 11 | 1.14.E-08 | 2 | 0.654 |
| mitochondrial electron transport, cytochrome c to oxygen | 24 | 6 | 3.01.E-04 | 1 | 0.683 |
| mitochondrial electron transport, NADH to ubiquinone | 53 | 11 | 1.69.E-07 | 2 | 0.697 |
| proton motive force-driven mitochondrial ATP synthesis | 68 | 14 | 1.02.E-09 | 7 | 0.015 |
| oxidoreduction-driven active transmembrane transporter activity | 78 | 16 | 3.33.E-11 | 4 | 0.780 |
| proton-transporting ATP synthase activity, rotational mechanism | 20 | 4 | 3.09.E-02 | 3 | 0.280 |
| aerobic electron transport chain | 96 | 19 | 3.64.E-13 | 4 | 1.000 |
| mitochondrial ATP synthesis coupled electron transport | 104 | 19 | 1.71.E-12 | 4 | 0.844 |
| oxidative phosphorylation | 150 | 24 | 1.22.E-14 | 9 | 0.086 |
| ATP biosynthetic process | 107 | 15 | 5.53.E-08 | 8 | 0.043 |
| aerobic respiration | 202 | 27 | 1.50.E-14 | 10 | 0.189 |
| mitochondrial respiratory chain complex assembly | 96 | 11 | 9.11.E-05 | 6 | 0.350 |
| nucleoside triphosphate biosynthetic process | 140 | 16 | 3.04.E-07 | 8 | 0.204 |
| primary active transmembrane transporter activity | 195 | 18 | 9.07.E-07 | 5 | 0.395 |
| purine ribonucleoside triphosphate metabolic process | 260 | 21 | 5.59.E-07 | 11 | 0.379 |
| nucleoside triphosphate metabolic process | 286 | 22 | 5.80.E-07 | 11 | 0.853 |
| proton transmembrane transport | 170 | 11 | 1.67.E-02 | 6 | 1.000 |
| nucleotide biosynthetic process | 335 | 21 | 1.64.E-04 | 10 | 0.940 |
| ribonucleotide metabolic process | 519 | 26 | 1.22.E-04 | 14 | 0.868 |

(e)

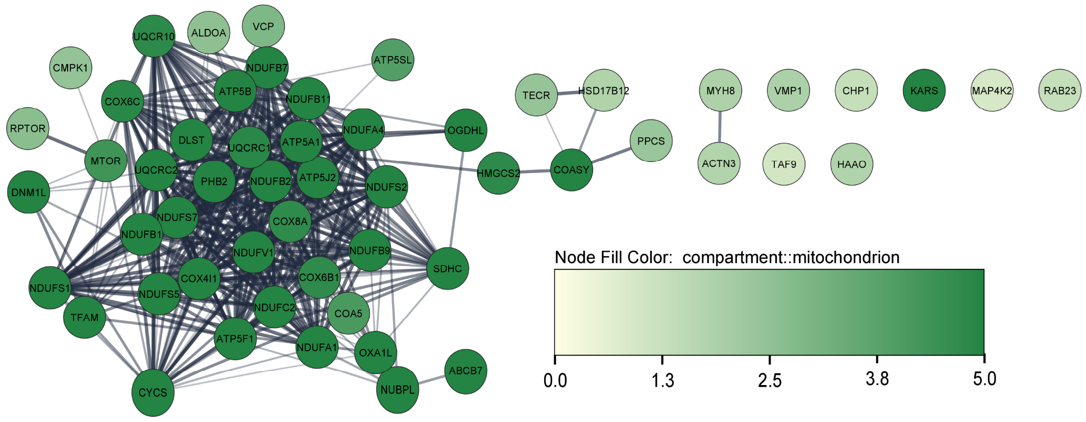


**Supplementary Figure 14 |** Detailed analysis of OxPhos terms and their annotated genes. (a) The number of genes in each region of Fig. 4a separated by -1.65 and 1.65 of z-RE for both x- and y-axis. Percentages are shown in parentheses underneath each gene number. (b) Pair-wise plot of z-RE obtained from CD63/CD9-CIBER screening for 722 genes annotated to OxPhos terms. Dashed lines show +/-1.65. (c) The number of genes in each region of Supplementary Fig. 14b. OxPhos genes are concentrated to CD63-CIBER-specific lower hits (middle left region; region 2) compared to all the screened genes (p = 0.00012: Fisher's exact test). (d) Table of OxPhos terms significant for lower hits of CD63-CIBER. "Term size" shows the number of genes annotated to a term in each row. "#Associated genes" shows the number of genes annotated to the term and detected as hits in each screening. adjp; p values by Fisher's exact test adjusted by Bonferroni step down. (e) STRING analysis of OxPhos genes on Fig. 4b (z-RE lower than -1.65 in either screen). Node color corresponds to mitochondrial associations according to Cytoscape. The density of the edge is proportional to the strength of PPI.

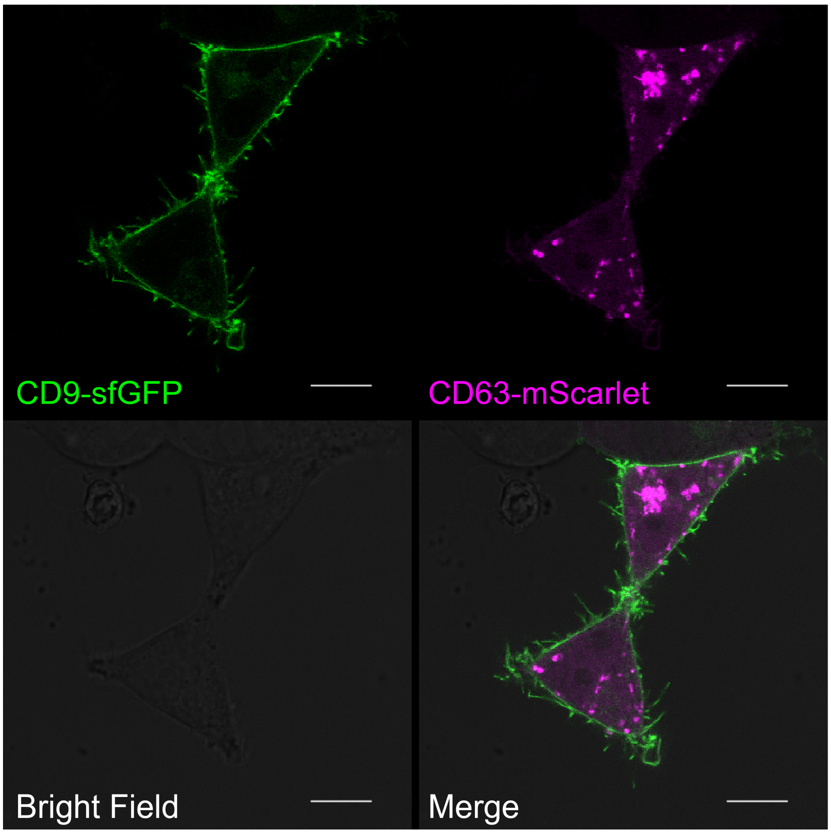


**Supplementary Figure 15 |** Subcellular localization study of CD63 and CD9. HEK293T cells seeded on an 8-well chamber were transiently co-transfected with CD9-sfGFP and CD63-mScarlet expression vectors. After a day of incubation, cells were visualized with a Leica SP8. Fluorescence images were captured with excitation and emission wavelengths of 488/510-555 nm for sfGFP and 569/579-665 nm for mScarlet. Scale bar, 10 µm.

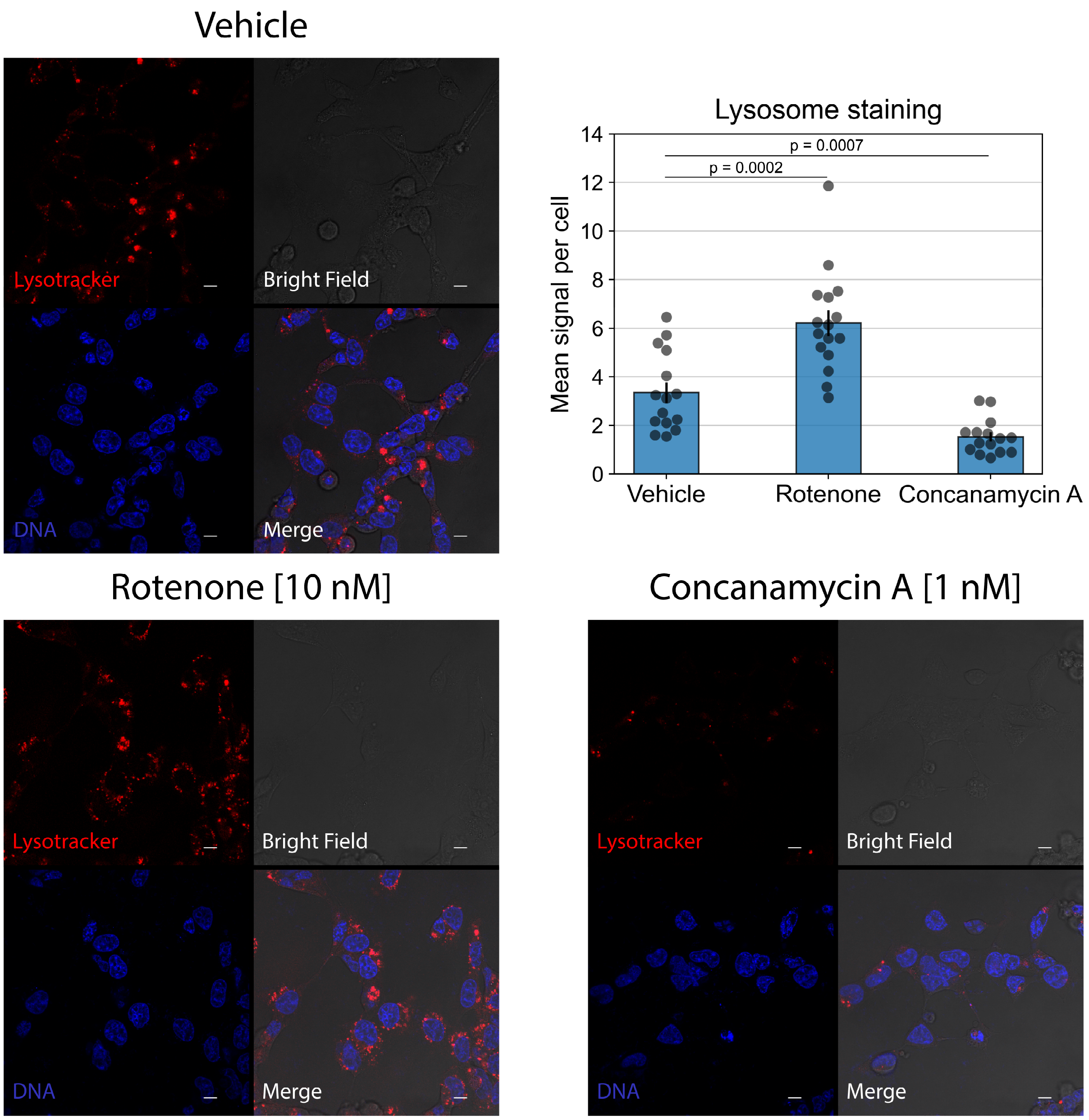


**Supplementary Figure 16 |** Confirmation of lysosomal activity upon treatment with inhibitors. HEK293T cells seeded on an 8-well chamber were treated with rotenone at 10 nM or concanamycin A at 1 nM for 24 hours. After the treatment, cells were stained with Hoechst33342 and then LysoTracker™ Red DND-99. Cells were visualized with a Leica SP8. Fluorescence images were captured with excitation and emission wavelengths of 577/584-718 nm for LysoTracker™ Red DND-99 and 405/440-510 nm for Hoechst33342. Scale bar, 10 µm. The boxplot at the upper left shows the mean LysoTracker™ signal per cell in the displayed image for each condition. Error bars represent ± SEM of fluorescence signal from 15~16 different cells in a single experiment. p: two-tailed Welch's t-test.

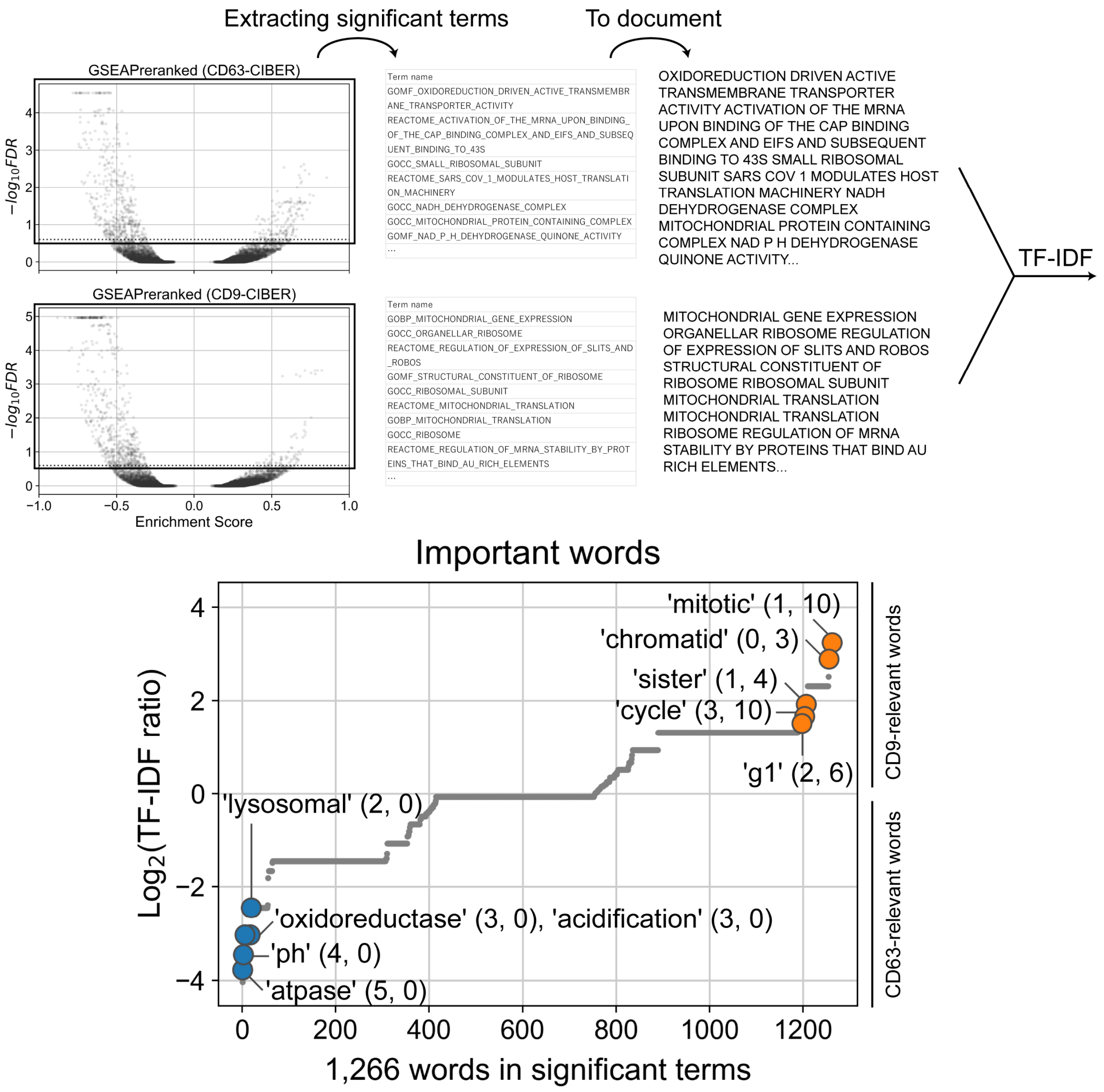


**Supplementary Figure 17 |** Term frequency - inverse document frequency (TF-IDF) analysis of hit terms. Words to be analyzed were collected from terms with FDR lower than 0.25 in the results of GSEAPreranked of CD63-CIBER and CD9-CIBER screening. The figure shows the $\log_2$ ratio of TF-IDF scores (CD9 vs CD63) from 1,266 words detected. Annotations were expressed as ‘word’ (count in CD63-CIBER, count in CD9-CIBER). This analysis suggests that the mitotic events are more relevant to $CD9^+$ sEVs (‘sister’ is used as SISTER_CHROMATID or SISTER_CHROMATIDS in terms, and 8 out of 10 ‘cycle’ were used as CELL_CYCLE in terms). Note that the $CD63^+$ EV-specific involvement in lysosomal activity is again detected in this analysis. See Supplementary Data 8 for individual TF-IDF scores.

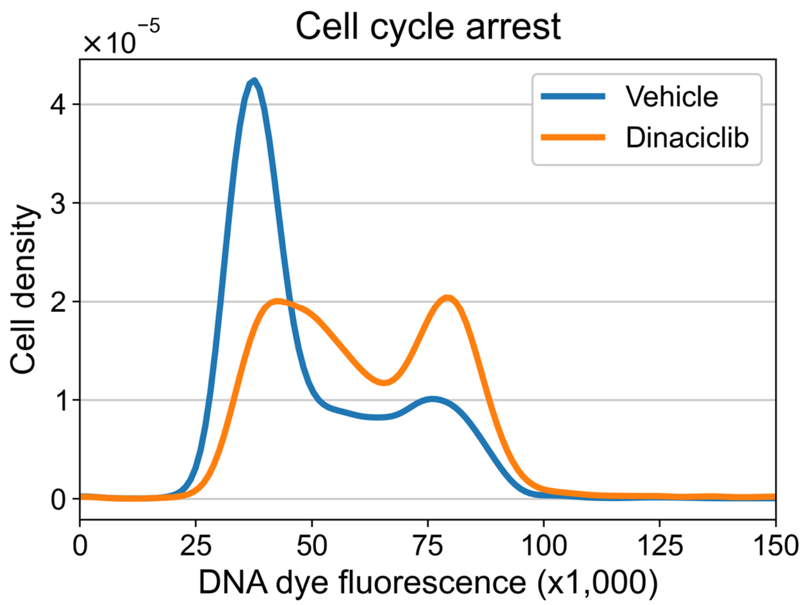


**Supplementary Figure 18 |** Flow cytometry analysis of cell cycle arrest by dinaciclib. HEK293T cells were treated with dinaciclib at 10 nM for 24 hours, stained with a DNA dye and analyzed by FCM. Curves for each condition shows kernel density estimation calculated from the FCM histogram. This data shows that the cells were successfully arrested at G2/M phase upon treatment with dinaciclib.

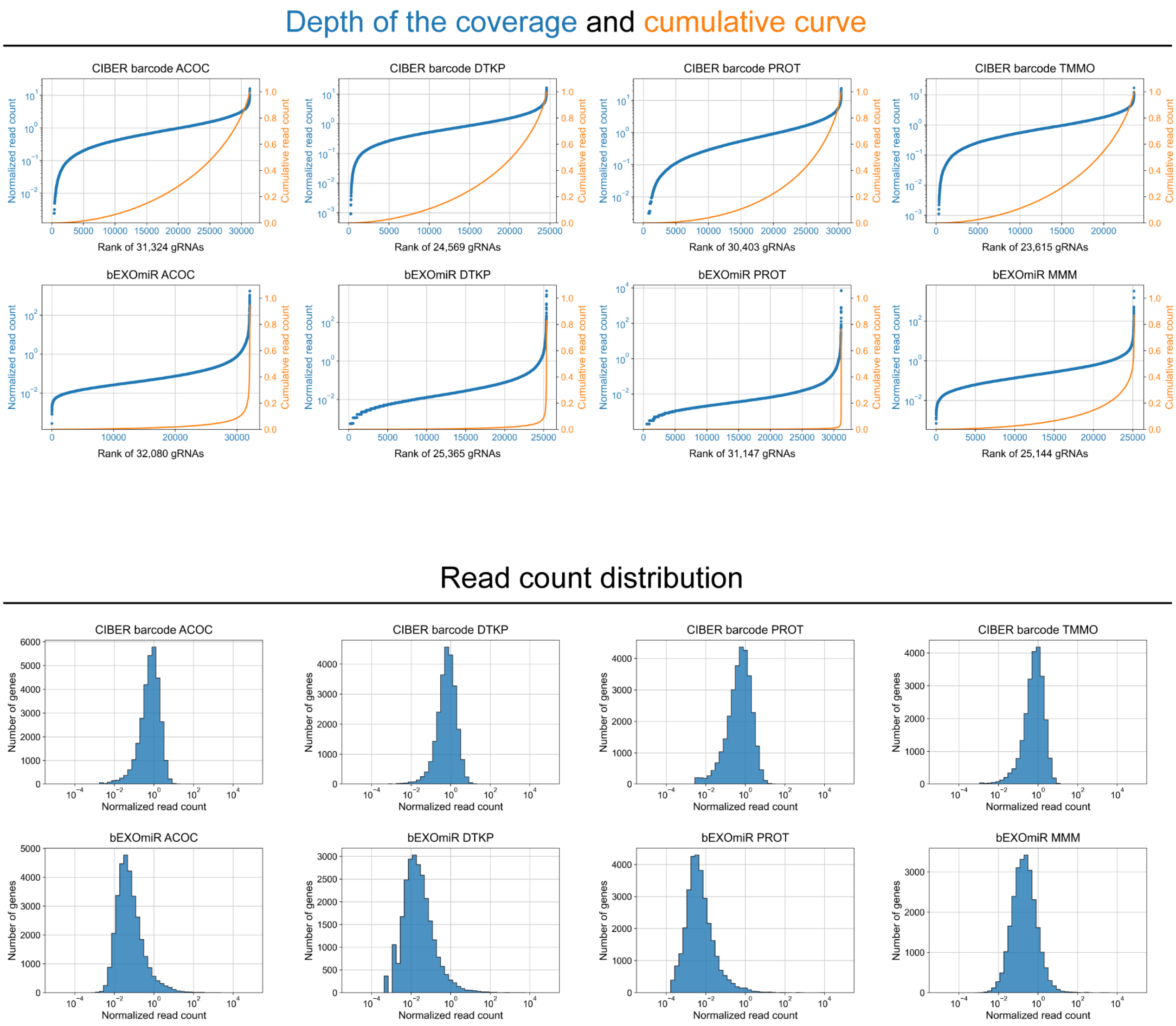


**Supplementary Figure 19 |** Comparison of sequencing results of CIBER screening and a previous report[15]. The upper panel shows the depth of barcode coverage from sEVs released by Cas9-expressing cells (the data of CIBER barcode ACOC/DTKP/PROT/TMMO is the same as shown in Supplementary Fig 6(b)). In the CIBER-screening barcodes were uniformly detected, whereas most reads covered a limited population of barcodes in the previous report. The lower panel shows the histogram with the distribution of barcodes by read count. Barcodes are symmetrically distributed in CIBER screening while they are right-skewed in the previous report. These data support the superior reliability of sEV barcoding in the CIBER screening platform.

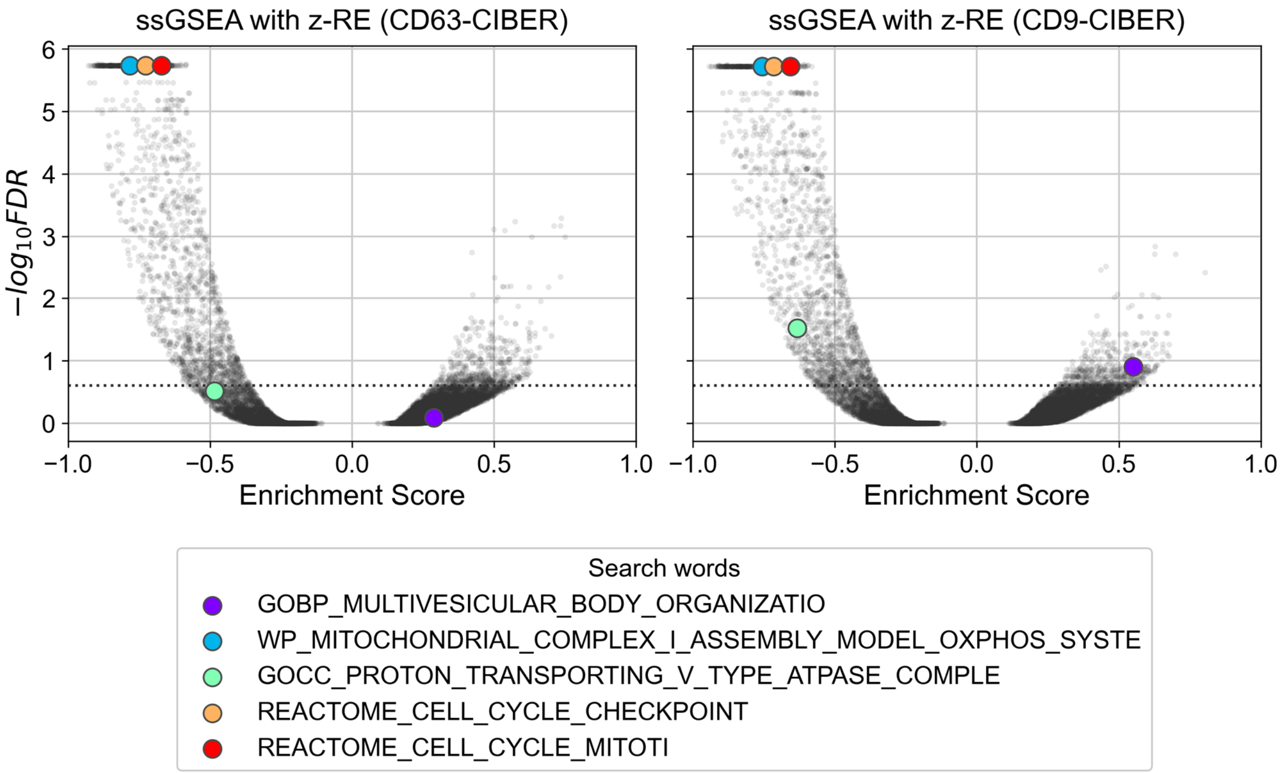


**Supplementary Figure 20 |** Volcano plot of enrichment score calculated via GSEAPreranked with z-$FC_{sEVs}$ as the query array. The dashed line shows FDR of 0.25. Because $FC_{sEVs}$ are highly correlated to $FC_{cells}$ (Fig. 2b), genes affecting cellular activity (e.g. viability) are often detected as false-positive lower hits in the screening when only $FC_{sEVs}$ are used to find hit genes. It is also impossible to see the different effect of the cell cycle on the release of $CD63^{+}$ sEVs and $CD9^{+}$ sEVs when z-$FC_{sEVs}$ are used as a pre-ranked query array.

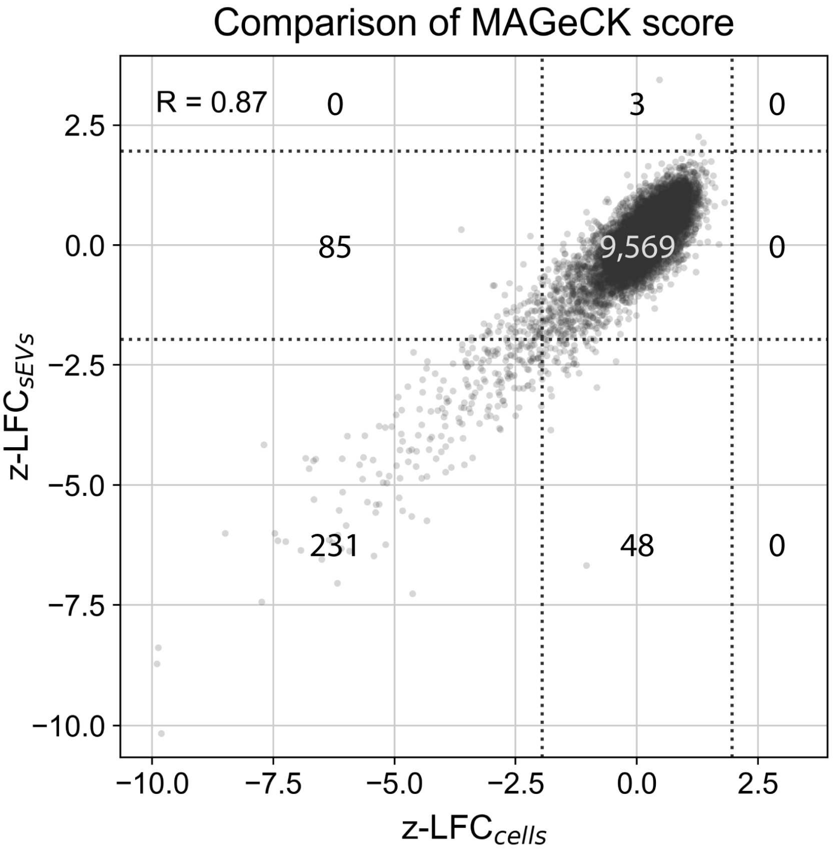


**Supplementary Figure 21** | Data showing the applicability of the sEV-barcoding system for cell-free CRISPR screening. The pair-wise plot of z-normalized $\log_2$ fold-change of the MAGeCK scores[16] of 9,936 genes (z-LFC, Cas9+ vs Cas9-) are shown. The scores are calculated from gRNA abundance in sEVs and cells with the data of CD63-CIBER screening (excluding the hit genes of the CD63-CIBER screening; i.e. CIBER screening was used as the counter assay). The numbers in each region divided by dashed lines show the number of genes in it. z-$LFC_{cells}$ is almost equivalent to the output of normal CRISPR screening, reflecting the changes of the cell population resulting from gene KO. Three hundred and sixteen genes were detected as lower hits under a threshold of +/- 1.96, an empirical threshold for selecting the top/bottom 5% of the population. z-$LFC_{cells}$ and z-$LFC_{sEVs}$ were highly correlated (R = 0.87) and the cell-free output obtained by sequencing barcodes in sEVs (z-$LFC_{sEVs}$) successfully detected 231 genes out of 316 hit genes calculated from the cellular barcodes. Dashed lines show +/-1.96.

### Supplementary methods

#### Creation of the MS2-gRNA library

The MS2-gRNA library was created by replacing the scaffold region of the hCRISPRa-v2 h6 library (addgene #83985, 13,145 gRNAs) with the MS2-gRNA scaffold in pKK49. Three µg of hCRISPRa-v2 h6 library and 5 µg of pKK49 were digested with BlpI/NheI at 37°C for 3 hours. After agarose gel purification, each fragment was eluted with 24 µL of MilliQ. Five µL of spacer-containing fragments and 8 µL of scaffold-containing fragments were ligated with 13 µL of Ligation high Ver.2 (cat. # LGK-201, TOYOBO) at 16°C overnight. After EtOH precipitation, the ligated product was amplified following a protocol provided by Joung et al[17].

The diversity of gRNA in the created library was quantified by NGS. The spacer regions were amplified by PCR in a 50 µL reaction mixture composed of 45 µL of Platinum PCR SuperMix High Fidelity (cat. #12532016, Invitrogen), 4 µL of the library [35 ng/µL], 0.5 µL of Oligo #5 [5 µM] and 0.5 µL of Oligo #18 [5 µM]. PCR conditions were as follows: 94°C for 2 minutes, 40 cycles of (94°C for 30 seconds, 65°C for 30 seconds, 68°C for 30 seconds), 68°C for 30 seconds and 4°C hold. The PCR product was purified using AMPure XP according to the manufacturer's protocol and eluted in 20 µL of nuclease-free water. Sequencing and downstream analysis were performed as described in the main methods.

#### Flow cytometry analysis of knockout efficiency

The gRNA-transferring virus were prepared with gCtrl (pKK90) or gBFP (pKK209) on a well of a 12-well plate as described in the main methods. Wild-type HEK293T cells, HEK293T cells expressing either of Cas9 or CD63-dCas9 and HEK293T cells expressing both Cas9 and CD63-dCas9 were transduced with the virus and selected on puromycin at 0.3 µg/mL for 1 week. Cells were subjected to flow cytometry (FCM) analysis on a BD LSR II (Becton, Dickinson and Company) to measure the BFP intensity using the Pacific Blue channel. FCM data was processed using FACS Diva software (ver. 4.1).

#### Quantitative polymerase chain reaction (qPCR)

Transfection was performed on a 96-well plate at 1/16 scale of the 12-well plate format described in the main methods related to Fig. 3f. Cells were cultured for 24 hours and then lysed using a SuperPrep II Cell Lysis Kit for qPCR (cat. #SCQ-501, Toyobo) according to manufacturer's protocol. Two µL of lysate was applied for 1 step qPCR using RNA-direct SYBR Green Realtime PCR Master Mix (cat. #QRT-201, Toyobo. Twenty µL of reaction mixture consisted of 2 µL of lysate, 10 µL of Master Mix, 1 µL of $Mn(OAc)_2$ at 50 mM, 0.1 µL of forward primer at 50 µM, 0.1 µL of reverse primer at 50 µM and NFW up to 20 µL). PCR conditions were as follows; 90 °C for 30 seconds, 61°C for 20 minutes, 95°C for 30 seconds, 45 cycles of (95°C for 5 seconds, 55°C for 10 seconds and 74°C for 15 seconds). Relative mRNA level was calculated using the $2^{-\Delta\Delta Ct}$ method with GAPDH as an internal control. Primer sequences are listed in Supplementary Table 2.

**Localization study of CD63 and CD9**

40,000 HEK293T cells in 200 µL of culture medium were reverse-transfected with 100 ng of pKK151 (CD9-sfGFP) and 100 ng of pRK397 (CD63-mScarlet) on an 8-well chamber (cat. #ib80826, Ibidi). After 24 hours of incubation, cells were visualization by a confocal fluorescence microscope (TCS SP8, Leica). The excitation and emission wavelengths can be found in the figure legends.

**Visualization of cellular lysosomal activity**

Wells of an 8-well chamber were coated with 200 µL of collagen solution (cat. #TMTCC-050, Toyobo) for 1.5 hours at room temperature and washed with 200 µL of PBS twice. HEK293T cells were plated ($3.0 \times 10^4$ cells in 200 µL of supplemented media) on the coated wells and allowed to expand to around 70% confluency. Media were changed to 200 µL of optiMEM containing either of vehicle (0.1% DMSO), rotenone at 10 nM or concanamycin A at 1 nM. After 24 hours of incubation, cells were stained with Hoechst33342 (Invitrogen, H1399) at 10 µg/mL for 10 minutes in DPBS and then LysoTracker™ Red DND-99 (Invitrogen, L7528) at 50 nM for 30 minutes in supplemented media before visualization by a confocal fluorescence microscope. The excitation and emission wavelengths can be found in the figure legends.

**TF-IDF**

Terms with FDR lower than 0.25 in the results of GSEAPreranked (Supplementary Data 6) were corrected and concatenated to prepare a document-like array. All the underscores in the array were replaced by spaces and the first words of each term showing ontology were removed. The TF-IDF scores for each word were calculated following the equation: $TF\text{-}IDF(t, d) = TF(t, d) \times (log_{10}(2/DF(t)) + 1)$, where $TF(t, d)$ indicates the number of the word $t$ in a document $d$ and $DF(t)$ indicates the number of documents that contains the word $t$. Any 0 scores are replaced by (minimal TF-IDF score in a document)/2 to avoid division by zero in calculating $\log_2$(TF-IDF ratio).

**Analysis for the application to cell-free CRISPR screening**

Raw read counts were analyzed to calculate z-LFC using MAGeCK software (ver 0.5.9.4, https://sourceforge.net/projects/mageck/files/0.5/mageck-0.5.9.4.tar.gz/download). The cellular counts from Cas9+ samples were used as the treated condition and that from Cas9- samples were used as the control condition to calculate $LFC_{cells}$. The sEVs counts were also processed similarly to calculate $LFC_{sEVs}$. The LFCs for each subpool library was z-normalized before being combined.
